# Structural basis for control of antibiotic production by bacterial hormones

**DOI:** 10.1101/2020.05.02.073981

**Authors:** Shanshan Zhou, Hussain Bhukya, Nicolas Malet, Peter J. Harrison, Dean Rea, Matthew J. Belousoff, Hariprasad Venugopal, Paulina K. Sydor, Kathryn M. Styles, Lijiang Song, Max J. Cryle, Lona M. Alkhalaf, Vilmos Fülöp, Gregory L. Challis, Christophe Corre

**Affiliations:** Department of Chemistry, University of Warwick, Coventry CV4 7AL, UK; School of Life Sciences, University of Warwick, Coventry CV4 7AL, UK; Warwick Integrative Synthetic Biology Centre, University of Warwick, Coventry CV4 7AL, UK; Department of Biochemistry and Molecular Biology, Monash University, Clayton VIC 3800, Australia; Infection and Immunity Program, Department of Microbiology, Monash University, Clayton, VIC 3800, Australia; Ramaciotti Centre for Electron Microscopy, Monash University, Clayton, VIC 3800, Australia

## Abstract

Actinobacteria produce numerous antibiotics and other specialised metabolites with important applications in medicine and agriculture. Diffusible hormones frequently control the production of such metabolites by binding TetR family transcriptional repressors (TFTRs), but the molecular basis for this remains unclear. The production of methylenomycin antibiotics in *Streptomyces coelicolor* A3(2) is initiated by binding of 2-alkyl-4-hydroxymethylfuran-3-carboxylic acid (AHFCA) hormones to the TFTR MmfR. Here, we report the X-ray crystal structure of an MmfR-AHFCA complex, establishing the structural basis for hormone recognition. We also elucidate the mechanism for DNA release upon hormone binding by single particle cryo-electron microscopy of an MmfR-operator complex. Electrophoretic mobility shift assays with MmfR mutants and synthetic AHFCA analogues illuminate the role played by individual amino acid residues and hormone functional groups in ligand recognition and DNA release. These findings will facilitate the exploitation of Actinobacterial hormones and their associated TFTRs in synthetic biology and novel antibiotic discovery.

## Introduction

Actinobacteria typically have a complex life cycle that begins with spore germination, leading to a dense network of branched multi-nucleoid hyphae^1^. Once nutrients become scarce, aerial hyphae begin to form. These septate, producing a series of mono-nucleoid compartments that differentiate to create chains of spores. The production of specialised metabolites is coordinated with this life cycle, usually commencing at the onset of aerial growth^2^.

Diffusible hormones are commonly employed to induce the expression of specialised metabolite biosynthetic gene clusters (BGCs) in Actinobacteria^3^. The archetypal example is A-factor, a γ-butyrolactone (GBL) that triggers the formation of aerial mycelium and the production of the antibiotic streptomycin in *Streptomyces griseus* (Figure 1)^4^. Binding of A-factor to ArpA, a TetR family transcriptional repressor (TFTR), releases it from the promoter of a transcriptional activator that induces the expression of genes controlling morphogenesis and antibiotic biosynthesis (Figure 1)^4^.

**Fig. 1.**
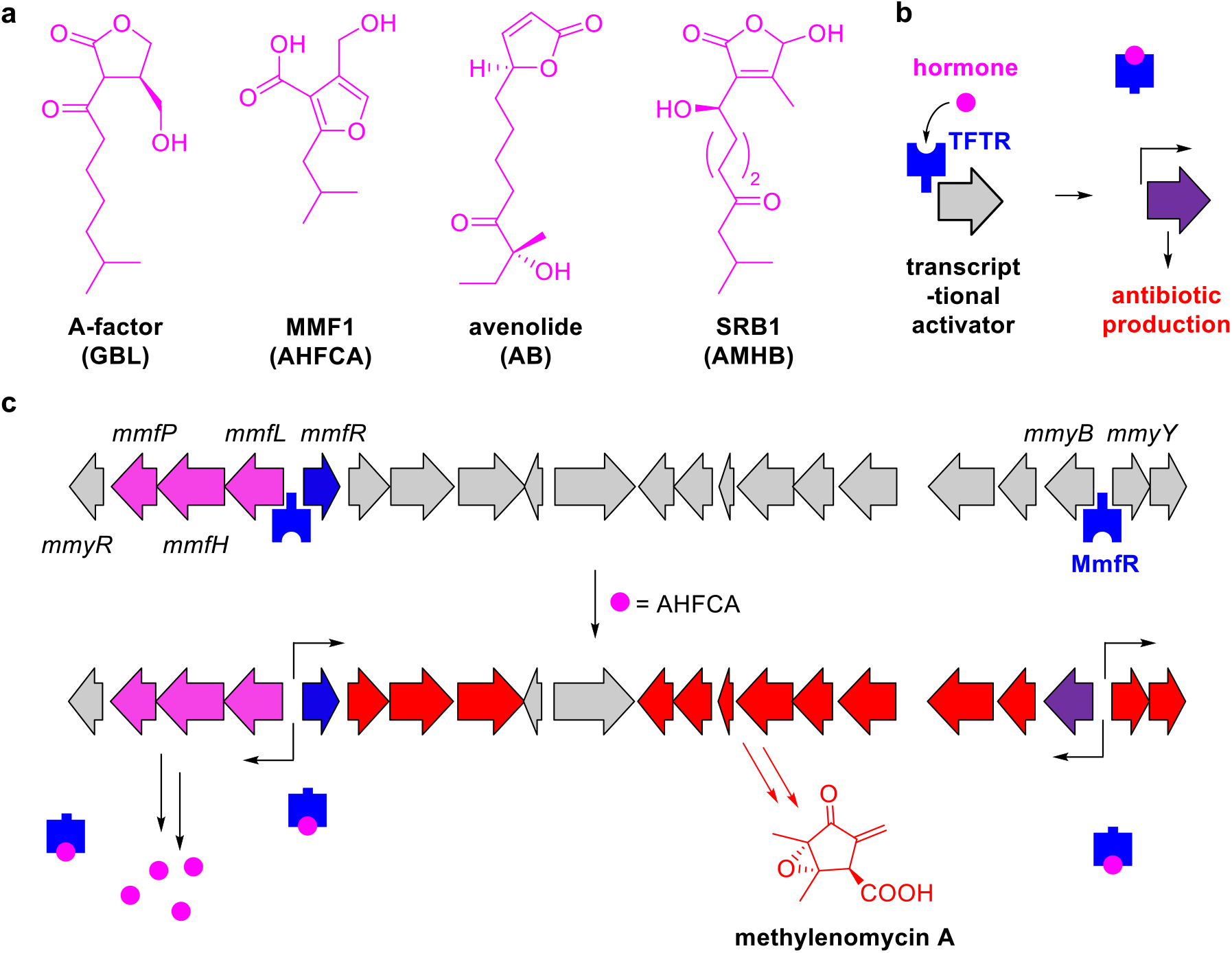
Classes of Actinobacterial hormone that induce antibiotic production by binding TFTRs and proposed mechanism for regulation of methylenomycin A biosynthesis by AHFCAs. **a**, Representative structures of the four hormone classes – γ-butryolactones (GBLs), 2-alkyl-4-hydroxymethyl-furan-3-carboxylic acids (AHFCAs), 4-alkylbutenolides (ABs) and 2-alkyl-3-methyl-4-hydroxybutenolides (AMHBs) – known to control antibiotic production in Actinobacteria. **b**, Generalised mechanism for induction of antibiotic biosynthesis, involving hormone-mediated de-repression of a transcriptional activator by a TFTR. **c**, Proposed mechanism for regulation of methylenomycin A biosynthesis in *S. coelicolor*. MmfR is a TFTR that represses the *mmfLHP* operon, in addition to *mmfR*, *mmyB* and *mmyY*. The AHFCA concentration steadily increases due to low-level expression of *mmfLHP*. Binding of the AHFCAs to MmfR upregulates *mmfLHP* expression, resulting in a feed forward loop. It also releases repression of the *mmyB* transcriptional activator of the methylenomycin biosynthetic genes.

Antibiotic production (and in some cases bacterial morphogenesis) is controlled by analogous mechanisms involving ArpA homologues in several other Actinobacteria, and for many years GBLs were believed to be only type of hormone involved. However, over the past decade three additional classes of hormone – 2-alkyl-4-hydroxymethylfuran-3-carboxylic acids (AHFCAs), 4-alkylbutenolides (ABs) and 2-alkyl-3-methyl-4-hydroxybutenolides (AMHBs) – have been implicated in the induction of antibiotic biosynthesis in Actinobacteria, via binding to ArpA-like TFTRs (Figure 1)^5–7^. Moreover, such TFTRs regulate the biosynthesis of several commercially important metabolites, including D-cycloserine, clavulanic acid, natamycin and ivermectin (all on the World Health Organization’s list of essential medicines), the streptogramins (used to treat vancomycin-resistant enterococcal infections), and tylosin (an antibiotic used in veterinary medicine)^8^. However, in most cases the hormones that these TFTRs respond to are unknown^9,10^.

Methylenomycin A is an antibiotic produced by the model Actinobacterium *Streptomyces coelicolor* A3(2) (Figure 1)^11^. We previously reported that a group of five AHFCAs called the methylenomycin furans (MMFs) induce the production of methylenomycin A in *S. coelicolor*^5^. A three-gene operon (*mmfLHP*) located at one end of the methylenomycin BGC directs MMF biosynthesis (Figure 1)^12^. The divergent *mmfR* gene directly upstream of this operon encodes an ArpA-like TFTR that is hypothesized to bind the *mmfR-mmfL*, and *mmyB*-*mmyY* intergenic regions (Figure 1)^12^. Binding of the MMFs to MmfR is proposed to release it from the *mmfR*-*mmfL*, and *mmyB*-*mmyY* intergenic regions, allowing *mmyB*, which encodes an activator of the methylenomycin biosynthetic genes, to be expressed^12^.

Although X-ray crystal structures of CprB (a putative GBL-binding TFTR in *S. coelicolor*) and a CprB-DNA complex have been reported, both the structural basis for hormone recognition and the mechanism of DNA release by ArpA-like TFTRs remain poorly understood^13–15^. Here we report structures of MmfR bound to an AHFCA and complexed with DNA duplex containing the operator from the *mmfL*-*mmfR* intergenic region, shedding light on hormone recognition and the mechanism of DNA release. We also report electrophoretic mobility shift assays (EMSAs) employing wild type and mutant MmfR proteins, as well as a synthetic library of naturally occurring AHFCAs and analogues, which illuminate the role played by key amino acid residues and hormone functional groups in ligand recognition and DNA release.

## Results and Discussion

### MmfR DNA binding and release by MMFs

We overproduced MmfR in *Escherichia coli* as a soluble N-terminal His_6_ fusion protein, enabling purification to homogeneity using nickel affinity chromatography (Supplementary Figure 1). ESI-Q-TOF-MS confirmed the identity of the purified protein (Supplementary Figure 1). EMSAs showed that MmfR binds the 194 base pair (bp) *mmfL-mmfR* intergenic region and a 100 bp internal fragment, in addition to the 230 bp *mmyB-mmyY* intergenic region and a 98 bp internal fragment (Figure 2). Competition experiments established that the protein binds preferentially to the *mmyB-mmyY* intergenic region (Figure 2). Bioinformatics analyses identified homologous 18 bp pseudo-palindromic operator sequences, hypothesised to be methylenomycin furan-autoregulator responsive elements (MAREs) in each of these intergenic regions^12^. EMSAs with DNA hairpins containing the 18 bp MARE1 and MARE2 sequences confirmed that MmfR binds to both of these (Figure 2). Increasing concentrations of MMF1 resulted in progressive release of the MmfR from each of the operators (Figure 2).

**Fig. 2.**
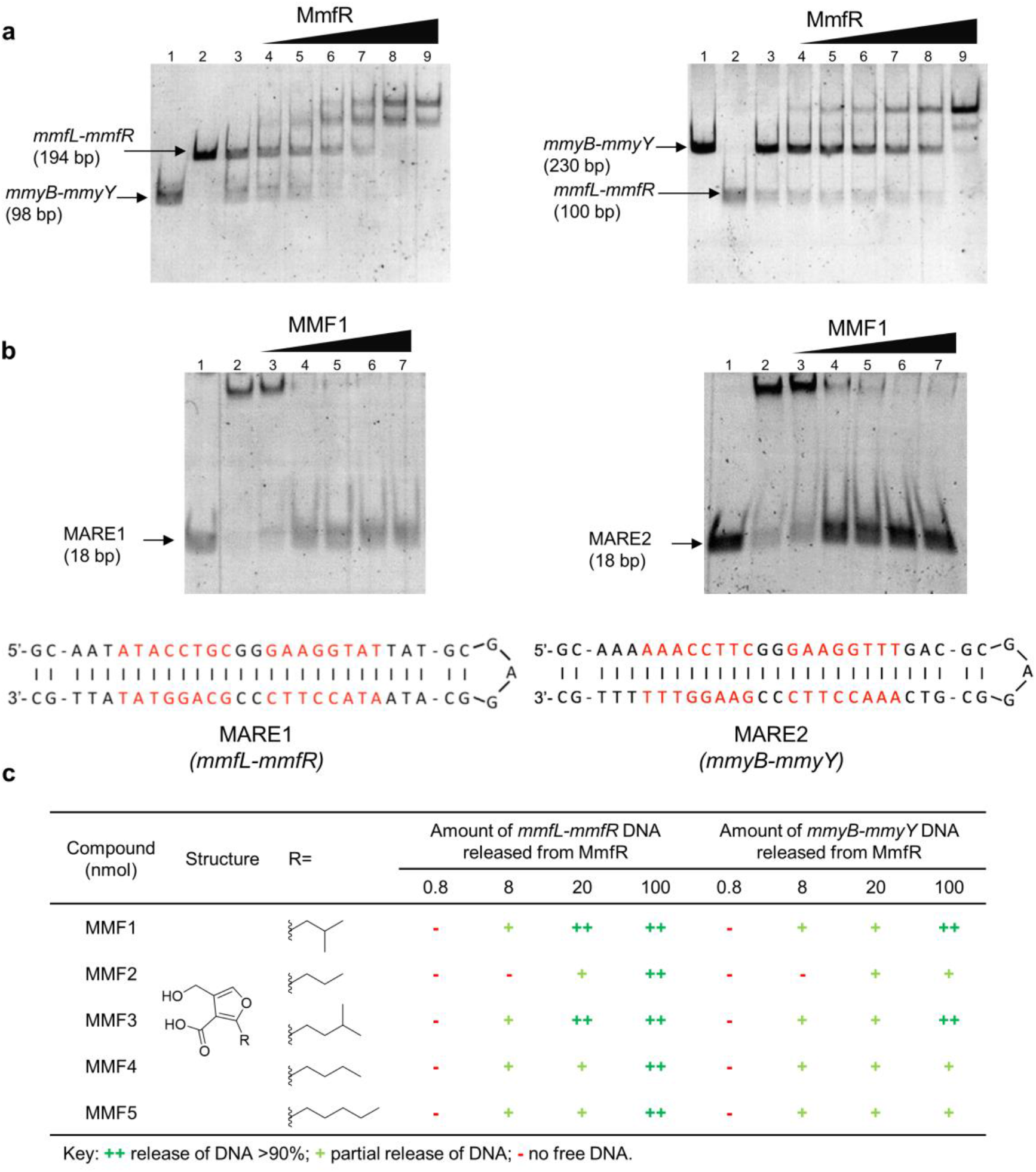
EMSAs showing that MmfR binds MAREs in the *mmfL*-*mmfR* and *mmyB*-*mmyY* intergenic regions and is released from the MAREs by MMF1. **a**, Competitive binding of MmfR to two pairs of DNA duplexes from the *mmfL-mmfR* and *mmyB-mmyY* intergenic regions. Lanes 1 and 2: individual DNA fragments (0.1 pmol); lane 3: equimolar mixtures of the two DNA duplex pairs (0.05 pmol of each); lanes 4 to 9: equimolar mixtures of the two DNA fragment pairs in the presence of increasing quantities of MmfR (0.1, 0.2, 0.3, 0.4, 0.5 and 0.6 pmol, respectively). **b**, Interaction of MmfR with short DNA hairpins containing MARE1 and MARE2, and progressive release of DNA from the complexes upon addition of increasing concentrations of MMF1. Lane 1: isolated DNA hairpins (0.8 pmol); lane 2: DNA hairpins mixed with MmfR (0.8 pmol and 4.0 pmol, respectively); lanes 3 to 7: addition of increasing quantities of MMF1 (0.8, 8, 20, 40 and 100 nmol respectively) to the DNA-protein complexes. **c**, Comparison of the quantities of the five naturally occurring MMFs required to release MmfR from the *mmfL-mmfR* (left) and *mmyB-mmyY* (right) intergenic regions.

The ability of each of the five MMFs produced by *S. coelicolor* to promote release of MmfR from the 194 bp *mmfL-mmfR* and 230 bp *mmyB-mmyY* intergenic regions was also established using EMSAs (Supplementary Figure 2). The amount of free DNA relative to the protein-DNA complex was estimated at four MMF concentrations (Figure 2). All of the MMFs induced dissociation of MmfR from both of the operators, but higher concentrations of MMF2, MMF4 and MMF5 were required to fully release MmfR from the *mmyB-mmyY* intergenic region than the *mmfL-mmfR* intergenic region. This is consistent with the higher affinity of MmfR for MARE2 than MARE1 (Figure 2). Subtle differences in the two operator sequences, in particular a terminal 5’-AAA…TTT-3’ sequence in MARE2 versus a 5’-ATA…TAT-3’ sequence in MARE1 may explain this observation.

We also investigated the minimum quantity of each MMF needed to trigger methylenomycin production in *S. coelicolor*, using a previously reported assay (Supplementary Figure 3 and Supplementary Table 1)^5^. Although all five hormones were able to induce methylenomycin production, MMF1 was found to be most active, whereas MMF2 and MMF3 were the least active; more than ten times the amount of MMF2 and MMF3 than MMF1 was required to trigger production of the antibiotic (Supplementary Table 1).

In addition to AHFCAs, *S. coelicolor* produces GBLs that directly control the expression of the coelimycin BGC by binding to the MmfR homologue ScbR, which represses the expression of the transcriptional activator KasO^16^. To investigate whether MmfR is specific for AHFCAs, or is also able to respond to other classes of endogenous hormone, we synthesised SCB1 (Supplementary Figure 4), one of the three most abundant GBLs produced by *S. coelicolor*. EMSAs showed that SCB1 is unable to induce dissociation of MmfR from the *mmfL-mmfR* intergenic region (Supplementary Figure 5), suggesting there is no crosstalk between the AHFCA and GBL-dependent regulation systems in *S. coelicolor*.

### Crystal structure of MmfR-MMF2 complex

MmfR readily crystallised and we were able to collect high quality X-ray diffraction data (Supplementary Table 2), but the structure could not be solved using molecular replacement. Thus, the selenomethione derivative of MmfR was prepared and crystallised. Collection of diffraction data for this derivative allowed us to solve and refine the structure to 1.5 Å resolution (Supplementary Table 2). We also co-crystallised MmfR with MMF2 and solved the structure of the complex at 1.5 Å resolution (Figure 3; Supplementary Figure 6).

**Fig. 3.**
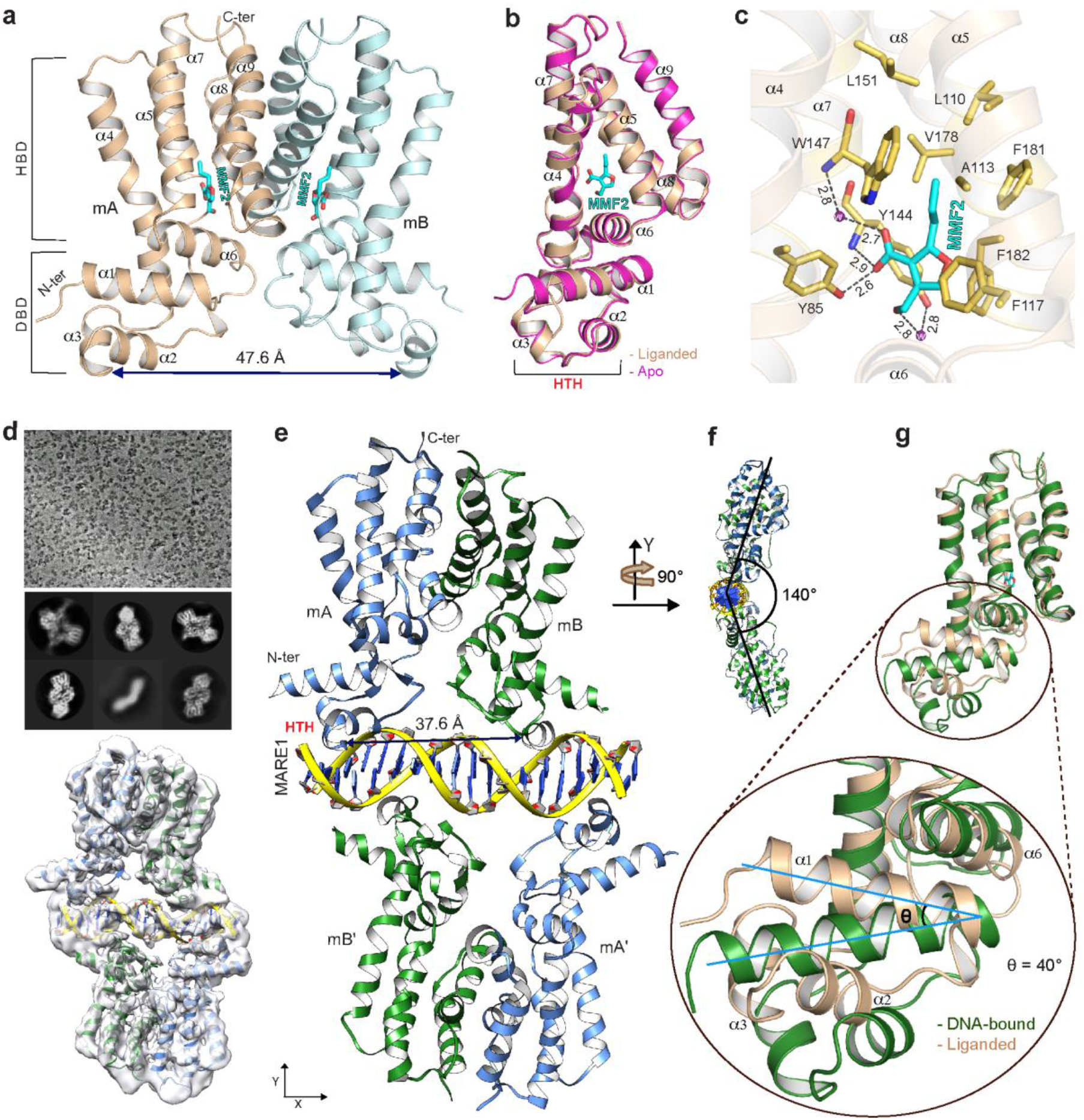
Structures of *apo-*MmfR, and the MmfR-MMF2 and MmfR-MARE1 complexes determined by X-ray crystallography and cryo-EM. **a,** X-ray crystal structure of the homodimeric MmfR-MMF2 complex; HBD = hormone-binding domain; DBD = DNA-binding domain. **b**, Overlay of the C_α_ atoms of *apo*-MmfR and the MmfR-MMF2 complex. **c**, Key residues lining the MmfR hormone-binding pocket, highlighting polar interactions (dashed lines) between MmfR (yellow) and MMF2 (cyan), in two cases mediated by ordered water molecules (purple spheres). All distances are in Å. **d**, Representative Cryo-EM image of the MmfR-MARE1 complex (top). Representative 2-D class averages of single particles (middle) and modelling of MmfR and the MARE1 DNA duplex into the cryo-EM density map using MDFF. **e**, Overall refined structure of the MmfR-MARE1 complex. Two homodimers of MmfR (monomer units in green and blue) bind to opposite faces of MARE1 (backbone in yellow and bases in blue). The monomer units in one of the MmfR homodimers are labelled mA and mB, whereas in the other homodimer they are labelled mA’ and mB’. HTH = DNA-binding helix-turn-helix. **f**, View of the MmfR-MARE1 complex rotated 90° about the y axis. **g**, Overlay of the HBD domain C_α_ atoms for MmfR in complex with the hormone (pink) and MARE1 (green), highlighting the different conformational state adopted by MmfR in the two complexes. The inset shows the axis of α-helix 1 tilts upwards by 40° in the MmfR-MMF2 complex relative to the MmfR-MARE1 complex, causing the HTH to pull away from the DNA major groove.

MmfR has a similar overall fold to CprB (Supplementary Figure 7), consisting of nine α-helices, among which the first three form the DNA-binding domain (DBD) containing the canonical helix-turn-helix (HTH)-motif in α-helices 2 and 3 (Figure 3)^13^. α-Helices 4-9 constitute the hormone-binding domain (HBD) and α-helices 8 and 9 form the homodimerisation interface (Figure 3), which contains primarily hydrophobic contacts. The *apo*-protein and the MmfR-MMF2 complex adopt very similar conformations (RMSD 0.437 Å for the 157 C_α_ atoms) (Figure 3).

Ten residues in the hormone-binding site interact directly with MMF2 (Figure 3). Six residues (L110, A113, W147, L151, V178, and F181) form a hydrophobic pocket that accommodates the alkyl chain. The F117 / Y144 and F182 side chains engage in parallel-displaced and T-shape π stacking interactions, respectively, with the furan. The carboxylate group of the hormone accepts hydrogen bonds from the Y85 hydroxyl group, the backbone N-H group of Y144 and an ordered water molecule, which also interacts with the backbone N-H group of W147. A second ordered water molecule is hydrogen bonded to the hydroxymethyl group of the hormone and the hydroxyl group of Y144.

### Cryo-EM structure of MmfR-DNA complex

Attempts to crystallize MmfR bound to DNA containing its operator sequences were unsuccessful. We thus employed single particle cryo-electron microscopy (cryo-EM) to elucidate the structure of the MmfR-DNA complex (Supplementary Table 3). The protein was complexed with a DNA duplex containing MARE1. Single particles of the protein-DNA complex were observed in cryo-EM movies (Figure 3). Two-dimensional classification of the particles showed they adopted several different orientations (Figure 3). Three-dimensional classification and subsequent refinement yielded a 4.2 Å density map containing clearly defined secondary structure elements (Figure 3 and Supplementary Figure 8). Superimposition of the MmfR X-ray crystal structure onto the map indicated that the conformation of the DBD changes upon DNA binding (Supplementary Figure 8). Thus, we performed molecular dynamics flexible fitting simulations to generate a model of the complex (Figure 3).

The model shows that two MmfR homodimers bind to opposite sides of the DNA duplex (Figure 3). The obtuse angle between the planes that bisect the monomers in each homodimer is 140° (Figure 3), consistent with that observed for other TFTRs that bind their operators as homodimeric pairs (Supplementary Figure 9)^14,15,17,18^. As in most other TFTR-DNA complexes, α-helix 2 and α-helix 3 of MmfR, encompassing the HTH-motif, serve as spacer and recognition helices, respectively. Comparison of the intra-dimer distance between DBDs (measured from the backbone nitrogen atoms of residue G64 in α-helix 3) showed that this decreases from 47.6 Å in the protein-hormone complex to 37.6 Å in the protein-DNA complex (Figure 3). The C_α_ atoms of the MmfR HBD in the hormone and DNA-bound states were superimposed to understand the conformational changes in the protein that cause it to be released from its operator upon hormone binding (Figure 3). This revealed an upward shift of the DBD towards the HBD in the protein-hormone complex (Figure 3), which prevents the HTH from binding in the major groove of the DNA duplex.

### Comparison with other ArpA-like TFTRs

To develop insight into the molecular basis for hormone recognition and the mechanism of signal transduction from the HBD to the DBD, we aligned the sequence of MmfR with other ArpA-like TFTRs of known ligand specificity (Figure 4)^19^. The level of residue conservation was mapped onto the structure of the MmfR-MMF2 complex (Figure 4).

**Fig. 4.**
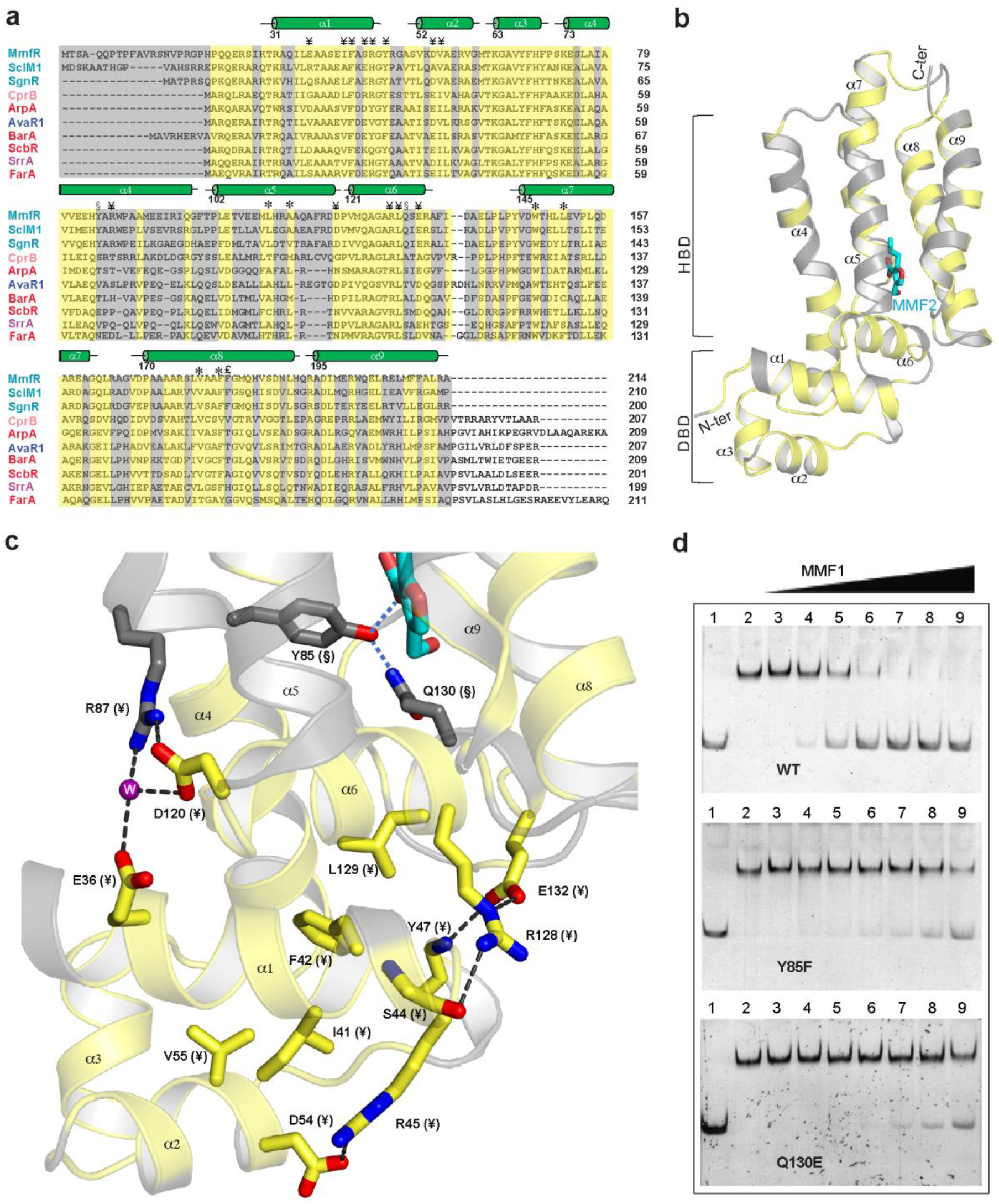
Insights into the mechanism of signal transduction in MmfR and other ArpA-like TFTRs. **a**, Multiple sequence alignment of TFTRs of known hormone specificity. Amino acids showing a high degree of conservation are coloured yellow, whereas those showing a low degree of conservation are coloured grey. Highly conserved residues hypothesised to be involved in the signal transmission from α-helices 4 and 6, through α-helix 1 to α-helices 2 and 3 in all TFTRs are marked ¥. The highly conserved residues and residue (F182), which is only conserved in AHFCA-binding TFTRs, lining the hydrophobic pocket of the HBD are indicated with * and £, respectively. The Y85 and Q130 residues, universally conserved in AHFCA-binding TFTRs, are marked §. Protein names are coloured according to the type of ligand each TFTR responds to; cyan: AHFCAs, red: GBLs, purple: AMHBs and blue: ABs. **b**, Mapping of residues showing a high (yellow) and low (grey) degree of conservation onto the structure of the MmfR-MMF2 complex. **c**, Close up view of the highly conserved residues (in yellow) proposed to mediate signal transmission from the HBD to the DBD in TFTRs via a network of hydrogen bonding and hydrophobic interactions. The Y85 and Q130 residues connect the carboxyl group of the ligand (in cyan) to α-helix 6 in AHFCA-binding TFTRs. Hydrogen bonding interactions and ordered water molecules are represented using dotted lines and purple spheres, respectively. **d**, EMSAs showing that Y85 and Q130 of MmfR play an important role in hormone-induced DNA release. Approximately ten times the quantity of MMF1 is required to release the Y85F and Q130E mutants of MmfR from the *mmfL*-*mmfR* intergenic region (194 bp) than the wild type protein. Lane 1: isolated DNA fragments (0.1 pmol); lane 2: DNA fragments mixed with MmfR (0.1 pmol and 1.8 pmol, respectively); lanes 3 to 9: addition of increasing quantities of MMF1 (0.8, 4, 8, 14, 20, 40 and 100 nmol respectively) to the DNA-protein complexes.

The W147, V178 and F181 residues, which form the sides of the hydrophobic pocket that accommodates the alkyl chain of the hormone in MmfR, are universally or very highly conserved in all other members of the ArpA family (Figure 4). On the other hand, while the three residues at the base of the pocket (L110, A113, L151) are highly conserved in AHFCA-binding TFTRs, they are less well conserved in proteins that bind other classes of hormone, although they are all still largely hydrophobic (Figure 4). This suggests that the alkyl chains common to the four known classes of hormone ligand for ArpA-like TFTRs (Figure 1) likely all bind in this hydrophobic pocket, with differences in the residues at the base of the pocket reflecting difference in the length and/or polarity of the alkyl chain.

The side chain NH_2_ group of Q130 in MmfR donates a hydrogen bond to the hydroxyl group of Y85, which is in direct contact with the carboxyl group of the AHFCA hormone (Figure 4). Q130 and Y85 are conserved in AHFCA-binding TFTRs, but not other members of the ArpA family (Figure 4). On the other hand, R128 and L129, positioned opposite Q130 on α-helix 6, are universally conserved in all ArpA-like TFTRs. The guanidinium group of R128 hydrogen bonds to the backbone carbonyl group of S44 and the side chain of L129 forms a hydrophobic contact with aromatic ring of the universally conserved F42 residue, which like S44 is located on α-helix 1 (Figure 4). Similarly, the carboxylate group of E132, which is also on the opposite face of α-helix 6 to Q130 and is very highly conserved in ArpA family members, hydrogen bonds to the backbone N-H and guanidinium groups of Y47 and R128, respectively (Figure 4). The side chains of two other very highly conserved residues, I41, located on the opposite face of α-helix 1 to F42, and V55, located on the top face of α-helix 2, also form a hydrophobic contact (Figure 4). Polar contacts between R45 in α-helix 1 and D54 in α-helix 2, and D120 at the N-terminus of α-helix 6 and E36 in α-helix 1 (via an ordered water molecule) also appear to be quite highly conserved in members of the ArpA family. This network of interactions suggests a plausible mechanism for signal transduction from the HBD to the DBD in MmfR. Binding of the carboxylate group of the hormone to the side chain of Y85 forces the C-terminal end of α-helix 6 to move downward and the N-terminal end of α-helix 4 to move inward (Figure 3). This pulls the N-terminal end of α-helix 1 towards the HBD, repositioning the HTH (Figure 3). It seems likely that other ArpA family members employ a similar signal transduction mechanism. However, Y85 and Q130 are typically T/V/V and S/T/A, respectively, in GBL, AB and AMHB-binding proteins, reflecting the structural differences between AHFCAs and these other hormone types.

To probe the involvement of Y85 and Q130 in hormone binding and signal transduction, we created Y85F and Q130E mutants of MmfR. The ability of the mutant proteins to bind the *mmfL*-*mmfR* intergenic region and for MMF1 to dissociate the resulting complexes was determined using EMSAs (Figure 4). In both cases, the mutant proteins bound tightly to the operator, but the concentration of MMF1 required to dissociate the protein from the DNA was approximately 10-fold higher than that required for the wild type protein. These results confirm that Y85 and Q130 play an important role in recognition of the hormone and transmission of the signal from the HBD to the DBD in AHFCA-binding TFTRs.

### AHFCA structure-activity relationship

The differences observed in the ability of MMF1-5 to dissociate MmfR from its operators indicate that the nature of the alkyl chain is one determinant of hormone recognition by AHFCA-binding TFTRs. To probe which structural features are important for hormone binding to MmfR further, a library of AHFCAs with variations in each of the furan substituents was synthesised (Figure 5). Structural alterations to the alkyl chain included shortening (analogues **1**-**4**), lengthening (analogues **5** and **6**), desaturation (analogue **7**), altering the position of the methyl branch (analogue **8**) and incorporation of an oxygen atom (analogue **9**). The carboxylic acid group was converted to the corresponding methyl ester (analogue **10**) and the hydroxymethyl group was replaced with a methyl group (analogue **11**) or a hydrogen atom (analogue **12**). The ability of these MMF analogues to promote dissociation of MmfR from the *mmfL-mmfR* intergenic region was assessed using EMSAs (Figure 5 and Supplementary Figure 10).

**Fig. 5.**
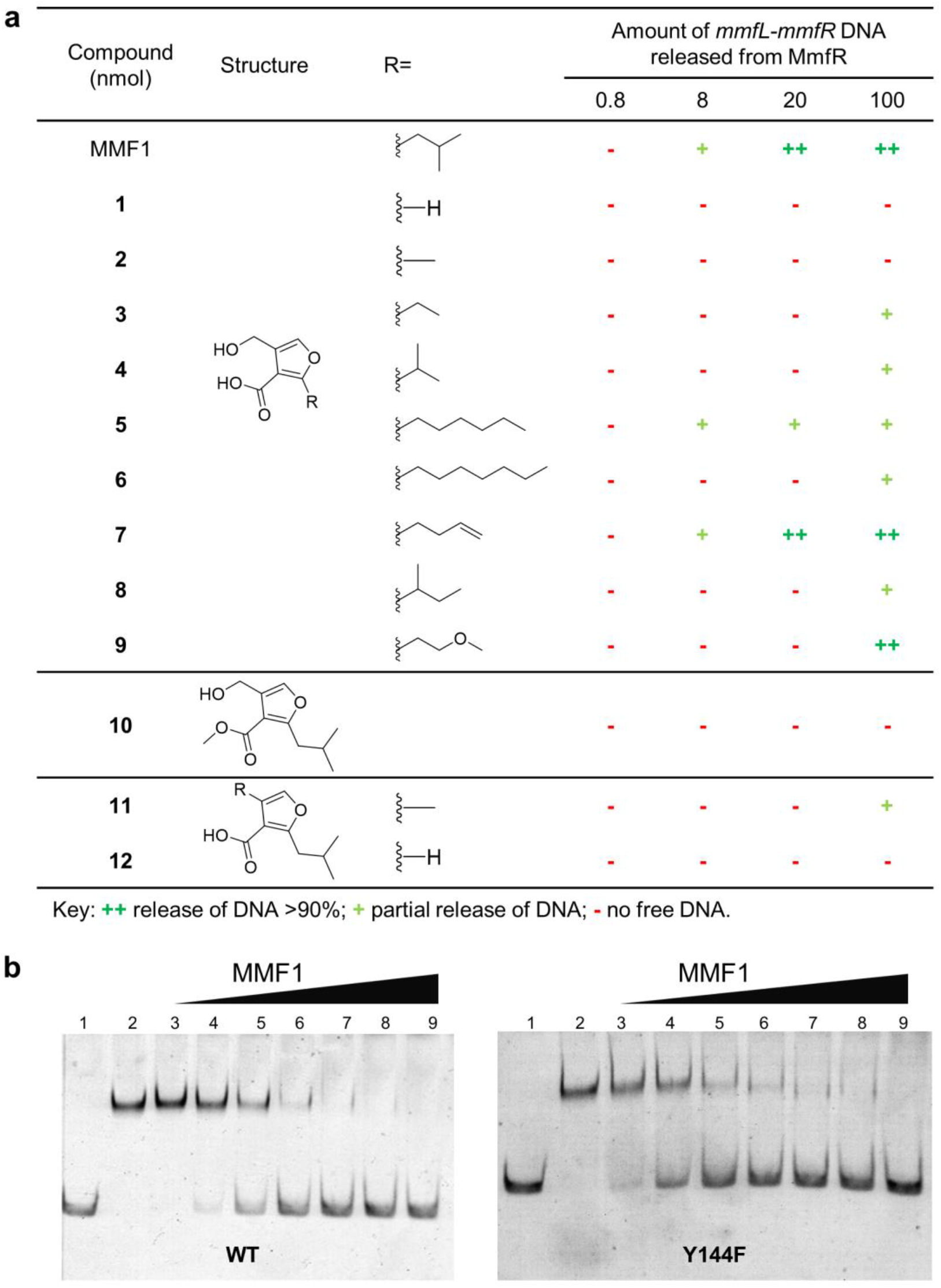
Probing the role of hormone functional groups in DNA release using an MMF analogue library. **a**, Results of EMSAs probing the ability of MMF analogues to release MmfR from the *mmfL*-*mmfR* intergenic region, relative to MMF1. **b**, EMSAs showing that similar quantities of MMF1 are required to release wild type MmfR and the Y144F mutant from the *mmfL*-*mmfR* intergenic region (194 bp). Lane 1: isolated DNA fragments (0.1 pmol); lane 2: DNA fragments mixed with MmfR (0.1 pmol and 1.8 pmol, respectively); lanes 3 to 9: addition of increasing quantities of MMF1 (0.8, 4, 8, 14, 20, 40 and 100 nmol respectively) to the DNA-protein complexes.

Moderate lengthening or desaturation of the alkyl chain does not appear to affect hormone binding to MmfR significantly. However, higher concentrations of analogues in which the alkyl chain had been more extensively lengthened or slightly shortened, or in which the methyl branch was repositioned or an oxygen atom was introduced, were required to effect full dissociation of MmfR from the operator. This effect was even more pronounced for analogues in which the alkyl chain had been extensively shortened (**2**) or completely removed (**1**). Similarly, removal of the hydroxyl group from the hydroxymethyl substituent (**11**) were modestly detrimental, whereas addition of a methyl group to the carboxyl substituent (**10**) and complete removal of the hydroxymethyl group (**12**) severely attenuated activity. However, it is surprising that the analogue lacking the hydroxyl group from the hydroxymethyl substituent still retains some activity, given that Y144 hydrogen bonds to the hydroxyl group via an ordered water molecule. To verify that the interaction between Y144 and the hydroxyl group of the hydroxymethyl substituent does not play an important role in hormone recognition by MmfR, we created a Y144F mutant. This mutant behaved very similarly to the wild type protein in EMSAs (Figure 5).

To determine whether the AHFCA structure-activity relationship we established *in vitro* is relevant to the induction of antibiotic biosynthesis *in vivo*, we investigated the ability of our AHFCA library to induce methylenomycin production using our previously described reporter system (Supplementary Table 4). The most active was analogue **7**, which is structurally similar to MMF4, but has a double bond at the extremity of the alkyl chain. Other compounds, with longer alkyl chains or altered methyl branch positions (e.g. **5**, **4** and **8**), in addition to analogue **11**, in which the hydroxymethyl group is replaced by a methyl group, showed a similar level of activity to MMF2 and MMF3. In contrast, **1**, **2** and **10** were all unable to induce methylenomycin production at quantities up to 50 μg. Similarly, SCB1 was unable to induce methylenomycin production, as expected from the results of the *in vitro* experiments.

## Conclusions

Despite the fact that A-factor, the first microbial hormone, and ArpA, its TFTR receptor, were discovered fifty and twenty five years ago, respectively, the molecular mechanisms by which they control gene expression has remained unclear^19,20^. The biosynthesis of several drugs on the World Health Organization’s list of essential medicines is controlled by TFTRs belonging to the ArpA subfamily. A detailed understanding of the molecular interactions between TFTRs, their DNA operators and the hormones that control them could be exploited to improve the production of such high value chemicals.

In this study, we have shown how binding of AHFCA hormones to ArpA-like TFTRs triggers antibiotic production in the model Actinobacterium *S. coelicolor*. By solving the crystal structure of MmfR in complex with one of its cognate hormones, MMF2, we have revealed the molecular basis for ligand recognition in AHFCA-binding TFTRs. In combination with our single particle cryo-EM structure of MmfR bound to the MARE1 operator, the crystal structure of the MmfR-MMF2 complex reveals a conserved mechanism for signal transduction in ArpA-like TFTRs.

The structures of only a handful of tetrameric TFTR-DNA complexes have been determined in the last two decades, highlighting the challenges associated with X-ray crystallographic analysis of such systems^21^. The MmfR-MARE1 structure is the first of a TFTR-DNA complex to be determined by single particle cryo-EM, which promises to significantly facilitate future efforts to illuminate the molecular mechanism of TFTRs.

## Methods

### Strains and plasmids

The strains and plasmids used in this study are summarized in Supplementary Table 5.

### Gene cloning

The *mmfR* gene was amplified from cosmid C73_787 by PCR using the primers listed in Supplementary Table 6. A CACC sequence was introduced at the 5’-end of the forward primer to allow for directional cloning of the blunt-ended PCR products into pET151/D-TOPO, resulting in the fusion of hexahistidine tag to the N-terminus of the recombinant protein. PCR products were purified using the GeneJET Gel Extraction kit (Thermo Scientific) and ligated with the linearized expression vector using the Champion pET151 Directional TOPO Expression kit (Invitrogen) according to the manufacturer’s guidelines. One Shot TOP10 chemically competent *E. coli cells* were transformed with the TOPO cloning reaction mixture. Transformants were selected on LB agar plates supplemented with ampicillin (100 μg/mL). Plasmids were purified from ampicillin-resistant colonies using the GeneJET Plasmid Miniprep kit and the integrity of the cloned gene was confirmed by sequencing (GATC Biotech).

### Protein overproduction and purification

For crystallization and EMSAs, recombinant His_6_-MmfR was overproduced in *E. coli* BL21 Star (DE3) cells carrying the pET151-*mmfR* plasmid. Cells were cultured in LB medium containing 100 μg/mL ampicillin at 37 °C to an OD_600_ of ~0.6. Expression was induced with 1 mM IPTG, and growth was continued at 20 °C overnight. Cells were harvested by centrifugation, resuspended in binding buffer A (20 mM HEPES, 0.5 M NaCl, 10% glycerol, 10 mM imidazole, pH 8), lysed by sonication, and clarified by centrifugation for 30 min at 4 °C. Cell lysates were passed through a 5 mL chelating sepharose FF column (GE Healthcare) charged with nickel and equilibrated in binding buffer. The column was washed with 100 mL binding buffer A, then 100 mL binding buffer containing 100 mM imidazole, and the protein was eluted in binding buffer containing 500 mM imidazole. Eluted protein was concentrated using a centrifugal concentrator with a 10,000 molecular weight cut-off (Vivaspin), and further purified by gel filtration chromatography on a Superdex S-200 column (GE Healthcare) equilibrated in buffer B (20 mM Tris-HCl, pH 8, 200 mM NaCl). Eluted protein was buffer-exchanged into buffer C (20 mM Tris-HCl, pH 8.0) and concentrated to 14 mg/mL using a centrifugal concentrator, frozen in aliquots of 100 μL in liquid nitrogen, and stored at −80 °C.

The same procedure was used to produce MmfR for cryo-EM analysis, except for the following. IPTG induction was carried out at 18 °C; the protein was purified on a Ni^2+^-nitrilotriacetic acid (Ni-NTA) sepharose resin (GE Healthcare), eluting with a linear gradient of imidazole (100 – 400 mM) in buffer A; and gel filtration was carried out on a HiPrep 16/60 Sephacryl S-200 HR column (GE Healthcare). The hexahistidine tag was removed by adding 1 mg/mL of purified recombinant His_6_-Tobacco Etch Virus protease to 10 mg/mL of MmfR and incubating at 20 °C for 12 h in buffer B containing 0.1 mM EDTA and 0.2 mM DTT^22^. The reaction mixture was passed through Ni-NTA resin equilibrated in buffer B to separate the His_6_-TEV protease from the cleaved MmfR protein. The flow through was further purified by gel filtration on a HiPrep 16/60 Sephacry S-200 HR column equilibrated in buffer B. The eluted protein was concentrated to 27 mg/mL using a 10,000 molecular weight cut-off centrifugal concentrator (Vivaspin).

### Protein crystallisation and X-ray data collection

Conditions for the crystallization of purified recombinant His_6_-MmfR were screened using a Honeybee crystallization robot. Sitting drops contained 200 nL of 14 mg/mL His_6_-MmfR and 200 nL reservoir solution. Reservoirs contained 75 μL of reservoir solution. Numerous hits were obtained and the best crystals grew in a solution containing 20% PEG 3,350 and 0.2 M of various divalent cations. Following optimization in hanging drops containing 1 μL protein and 1 μL reservoir solution, X-ray diffraction-quality crystals were grown in 10-15% PEG 3,350 and 0.2-0.25 M magnesium formate. Crystals were removed from sitting drops using a nylon loop, soaked briefly in LV cryo oil (MiTeGen), frozen and stored in liquid nitrogen. Crystals diffracted to 1.9 Å in-house using a sealed tube X-ray generator, and to 1.5 Å using synchrotron radiation at Diamond Light Source on beam line I24 (UK). Purified recombinant His_6_-MmfR (14 mg/mL) was mixed with 1 mM MMF2 and the resulting mixture was subjected to the crystallization conditions described above. These crystals also diffracted to 1.5 Å, using synchrotron radiation at Diamond Light Source on beam line IO4.

Selenomethionine-labelled protein was prepared by overproducing His_6_-MmfR in the methionine auxotroph *E. coli* B834 as described above, except minimal medium containing selenomethionine instead of methionine was used. The resulting protein was purified as described above, except 5 mM 2-mercaptoethanol was included in all purification buffers, and crystallized in several conditions, with the best crystals grown in a solution containing 8% PEG 8,000, 0.1 M magnesium acetate and 0.1 M sodium acetate, at pH 4.5. These crystals diffracted to 2 Å using synchrotron radiation at Diamond Light Source, and three complete datasets were collected at peak, inflection point and remote wavelengths at beam line IO3, following a wavelength scan.

### X-ray data processing, structure determination and refinement

All X-ray diffraction data were processed using XDS^23^. Further data handling was carried out using the CCP4 software package^24^. The structure was phased using only peak wavelength data from selenomethionine-labelled His_6_-MmfR crystals using a single-wavelength anomalous dispersion (SAD) approach. The SOLVE program located all seven selenated sites in the protein, and RESOLVE fitted 52% of the residues in the resulting electron density^25,26^. At this stage, model building was continued on the 1.5 Å data set from the isomorphous MmfR-MMF2 complex, and the model was further extended automatically by ARP and manually using O^27,28^. The unliganded structure was solved by molecular replacement using Phaser^29^. Refinement of both structures was performed using REFMAC5^30^. Water molecules were added to the atomic models automatically using ARP at the positions of large positive peaks in the difference electron density map, only where the resulting water molecule fell into an appropriate hydrogen bonding environment. The crystallographic asymmetric unit contains one subunit of the polypeptide chain in both structures. The structures of the dimers were generated from the crystallographic two-fold symmetry operators. The polypeptide could be unambiguously traced from residues 26-214 and 28-214 in the unliganded and MmfR-MMF2 complex structures, respectively. Data collection, phasing and refinement statistics are given in Supplementary Table 2.

### Cryo-Electron Microscopy

The protein-DNA complex was prepared by adding annealed MARE1 oligonucleotide (5’-ATACCTGCGGGAAGGTATT-3’) to 0.5-fold molar excess of cleaved MmfR and incubating at 20 °C for 1 h. It was purified by gel filtration using the column and protocol described above for cleaved MmfR and concentrated to 17.0 mg/mL. The MmfR-MARE1 complex was diluted to a concentration of 0.1 mg/mL in buffer B and the resulting solution was spotted onto plasma-cleaned Quantifoil carbon EM grids (hole size R1.2/1.3), which were plunge frozen in liquid ethane using an FEI Vitrobot Mark III (blotting time 3.0 s, 100% humidity, −3 mm blotting offset at 4 °C). Cryo-EM micrographs were collected using ThermoFisher Titan Krios™ equipped with a Gatan K2 Summit™ with Quantum-GIF energy filter operated at 300 kV at zero loss mode. A condenser aperture of 50 micron and FEI Voltage phase plate was used in the objective plane. Two Cryo-EM movie datasets (725 and 880 micrographs, each micrograph containing 50 frames) with 10 s exposure time and a total dose of 50 electrons per Å^2^ were recorded in counting mode at a nominal magnification of 165,000 (EFTEM mode), which corresponds to a calibrated pixel size of 0.84 Å. ThermoFisher Scientific EPU automated data collection software was used for data acquisition. Autofocus was set to keep the defocus at 0.5 micron and the phase plate position was advanced every hour.

The acquired movie frames were corrected for beam-induced translational motion and summed using MotionCor2 (version 2.1.10-cuda8.0) and the pixel size was raised to 1.092 Å by binning the dataset^31^. The summed images were saved and subjected to contrast transfer function (CTF) estimation employing GCTF (version 1.06-cuda8)^32^. Summed images with the best CTF fit, phase estimation and a defocus inside the range of 0.5–3.0 micron were selected manually by inspecting the thon rings fit to the theoretical resolution shells and these micrographs were retained for further image processing. This procedure resulted in datasets of 675 and 770 micrographs for MmfR-MARE1 complex. Subsequently, the micrographs were subjected to particle picking using crYOLO1.13^33^. One hundred images were chosen randomly and about 3,000 particles were picked manually to train the auto-picking module, which was then used to auto-pick particles from the CTF corrected and retained summed images of the complex. The trained module was able to pick 187,409 and 292,512 well-centred particles for the complex at a box size of 140 Å. The particles were then extracted and combined for reference-free 2-D classification in Relion3.0-beta to filter out low-quality particles^34^. This yielded a total of 379,033 particles, which were extracted at a box size of 180 pixels and a pixel size of 1.092 Å. The extracted particles were 3-D classified and an initial 3-D model was generated in Relion. From the 2-D and 3-D classes, it appeared that the complex has no symmetry. Thus, C1 symmetry was applied for all the model refinement steps and the particle size was not changed during data reduction and refinement. The particles resulting from tight masking and model refinement yielded an overall resolution of 4.2 Å with good angular distribution (Supplementary Figure 9).

### Modelling of the protein-DNA complex

The X-ray crystal structure of unliganded MmfR was manually aligned with the cryo-EM density map using Chimera^35^. MmfR was found to adopt four distinct positions in the map. B-form MARE1 DNA was also manually docked into the cryo-EM density map. The models were combined and the fit was optimised using molecular dynamics flexible fitting (MDFF)^36^. Runs were prepared with the VMD graphical user interface, and NAMD was used with the correction map and the CHARMM36 force field to perform the MDFF^37–40^. Figures were rendered in Chimera and PyMol (http://www.pymol.org).

### Electrophoretic mobility shift assays

The *mmfL-mmfR* (194 bp and 100 bp) and *mmyB-mmyY* (230 bp and 98 bp) intergenic DNA sequences containing the proposed binding sites for MmfR were amplified from cosmid C73_787 by PCR using the primers shown in Supplementary Table 6. The EMSAs with DNA fragments for the *mmfL-mmfR* or *mmyB-mmyY* intergenic regions were run on 6% non-denaturing polyacrylamide gels, while those with the short hairpin DNA fragments were run on 10% gels. For hairpin DNA, the synthetic single-stranded oligonucleotides (Sigma-Aldrich, Supplementary Table 6) were diluted in sterile water to 200 nM, warmed at 95 °C for 2 min and slowly cooled to promote hairpin formation^41^. In EMSAs with hairpin DNA, lane 1 contained 0.8 pmol of isolated DNA hairpin only and lanes 2-7 contained 0.8 pmol of hairpin DNA and 4.0 pmol of MmfR. For lanes 3 to 7, increasing concentrations of MMF1 were added (0.8, 8, 20, 40 and 100 nmol, respectively). For the competitive binding assays, each lane contained 0.1 pmol of DNA fragment(s) (individual or equimolar mixture) and for lanes 4 to 9, increasing quantities of MmfR was added. For other EMSAs, the amount of DNA and protein was kept constant at 0.1 pmol and 1.8 pmol, respectively. Compounds used in the EMSAs were dissolved in DMSO. The specific amount of each compound used was listed in the legend of the corresponding figure. For each 20 μL reaction, 4 μL of 5X binding buffer (100 mM HEPES, 50 mM (NH_4_)_2_SO_4_, 150 mM KCl, 5 mM EDTA, 5 mM DTT, 1% (w/v) Tween 20, pH 7.6) was used. DNA, protein, 5X binding buffer and distilled water were first mixed and incubated at room temperature for 15 min. After addition of compound solution, the resulting mixture was incubated at room temperature for 10 min, and 5 μL of loading buffer, containing 0.25X TBE, 34% (v/v) glycerol and 0.2% (w/v) bromophenol blue, were added. The total volume of each reaction did not exceed 25 μL. Samples were separated with a pre-run native polyacrylamide gel in 1X TBE buffer at 80 V at 4 °C until the loading dye had migrated to the bottom of the gel. On completion, the gel was incubated in a solution of GelRed (0.005% v/v) in 1X TBE buffer at room temperature for 30 min with agitation, then visualised on a UV transilluminator.

### Induction of methylenomycin production

As described previously, we added solutions of each compound at various concentrations directly to round plugs of AlaMM agar (pH 5.0, allowing optimal diffusion of methylenomycin) with 2-day old cultures of *S. coelicolor* W81 (MMF non-producer) growing on them^5^. Each compound was resuspended in methanol at diverse concentrations so that adding a 10 μL volume to the plug resulted in quantities of compounds ranging from 0.01 to 50 μg. The plugs were placed on an agar plate inoculated with *S. coelicolor* M145 (SCP1^−^, SCP2^−^, methylenomycin-sensitive) and, after 96-120 h of incubation at 30 °C, the extent of growth inhibition around the plugs (resulting from methylenomycin production) was recorded.

### Site-directed mutagenesis of MmfR

The Q130E, Y85F and Y144F mutants of His_6_-MmfR were created using the Q5 Site-Directed Mutagenesis Kit (New England Biolabs) according to the manufacturer’s instructions. The primers used are listed in Supplementary Table 6. All constructs were sequenced to confirm the presence of the desired mutations.

### Synthesis of MMFs and their analogues and SCB1

The procedures used (Supplementary Figure 11 and Supplementary Figure 12) and the compound characterisation data obtained are detailed in the Supplementary Information.

## Data availability

The atomic coordinates have been deposited in the Protein Data Bank for apo and liganded MmfR under the accession numbers 6SRM and 6SRN respectively. The three-dimensional cryo-EM map of protein-DNA complex has been deposited in the Electron Microscopy Data Bank (EMDB) under the accession number EMD-20781. This EMDB includes (1) raw half maps (2) B-factor sharpened map and (3) mask used for refinement and sharpening.

## Author contributions

S.Z., H.B., N.M., P.J.H., M.J.C., V.F., G.L.C. and C.C designed the experiments; S.Z., H.B., N.M, P.J.H., D.R., P.K.S., K.M.S., H.V., L.S. and C.C. executed the experiments; S.Z., H.B., N.M., P.J.H., D.R., M.J.B., H.V., L.J.S., M.J.C., V.F., G.L.C. and C.C. analysed data; and S.Z., H.B., L.A., G.L.C and C.C. wrote the manuscript.

## ACKNOWLEDGMENTS

C.C. acknowledges support of this work by a University Research Fellowship from The Royal Society (UF090255) and by grants BB/M022765/1 and BB/M017982/1 from the UK Biotechnology and Biological Sciences Research Council (BBSRC). G.L.C. is grateful to the Monash-Warwick Alliance (postdoctoral fellowship to H.B.), the University of Warwick (Chancellor’s International Scholarship to S.Z., Warwick Collaborative Postgraduate Research Scholarship to N.M. and Institute of Advanced Study Postdoctoral Research Fellowship to P.K.S.) and the ARC Centre of Excellence in Advanced Molecular Imaging for support of this work. Crystallographic data were collected at beam lines IO3, IO4 and I24 at Diamond Light Source, UK and we acknowledge the support of the beam line scientists.

