## Supplementary Information for "Structural basis for control of antibiotic production by bacterial hormones"

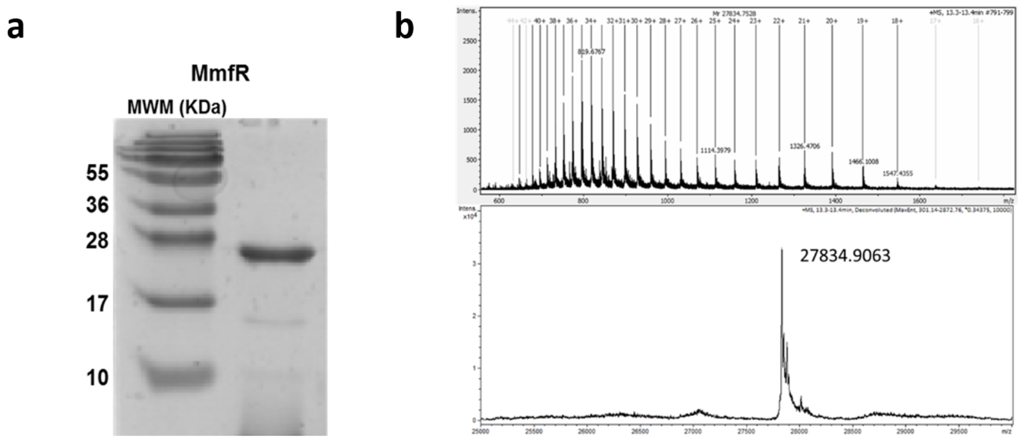

**Supplementary Fig. 1 | Analysis of purified recombinant His$_6$-MmfR.** **a**, SDS-PAGE. **b**, Mass spectrometry analysis: measured (top) and deconvoluted (bottom) mass spectra of His$_6$-MmfR (calculated mass = 27835.5 Da).

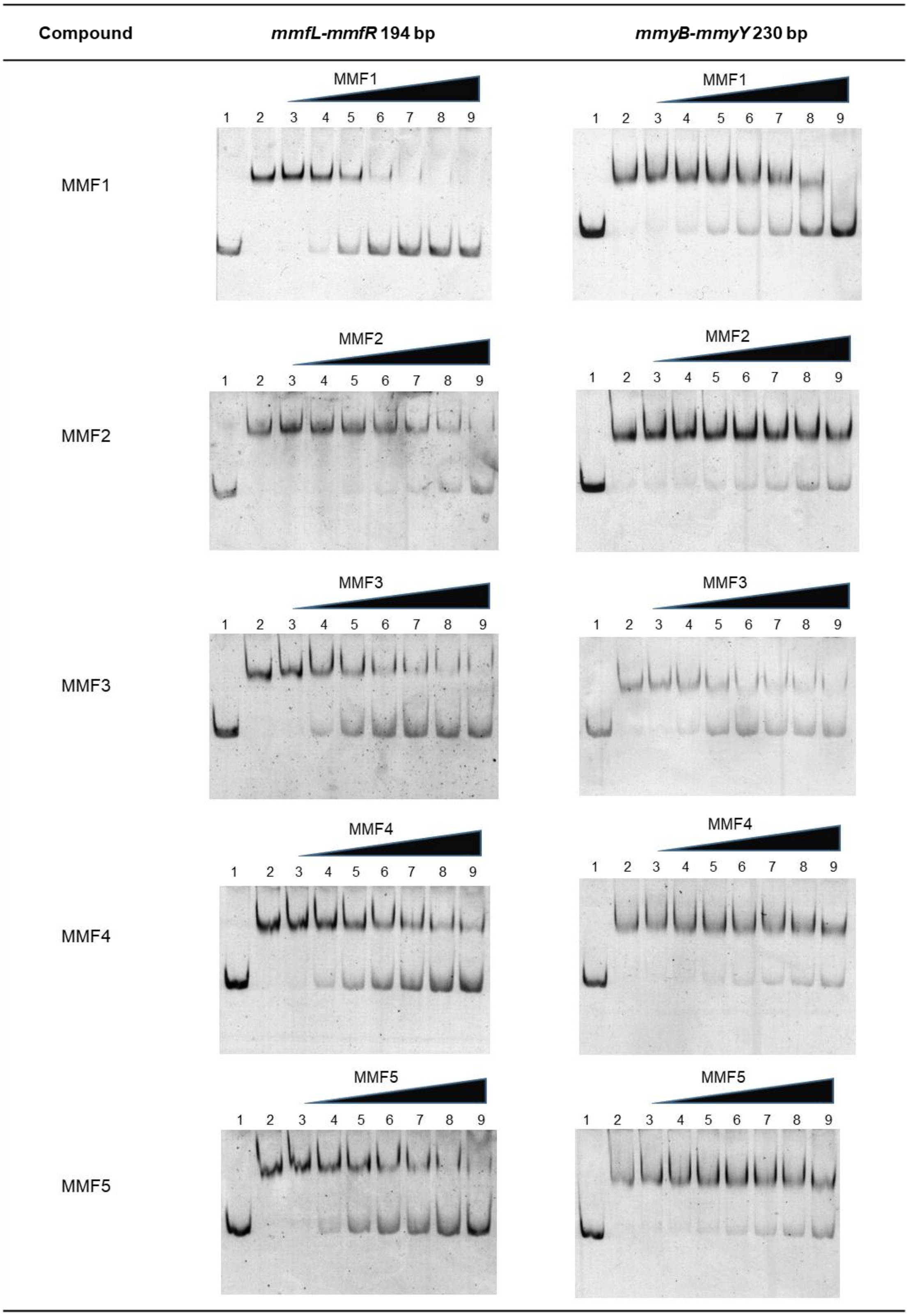

**Supplementary Fig. 2 | Interaction of MmfR with DNA fragments corresponding to the *mmfL-mmfR* and *mmyB-mmyY* intergenic regions (194 bp and 230 bp respectively) in response to increasing amounts of MMFs.** Lane 1: isolated DNA fragments (0.1 pmol); lane 2: DNA fragments mixed with MmfR (0.1 pmol and 1.8 pmol respectively); lanes 3 to 9: addition of increasing quantities of MMFs (0.8, 4, 8, 14, 20, 40 and 100 nmol respectively) to the DNA/protein complexes.

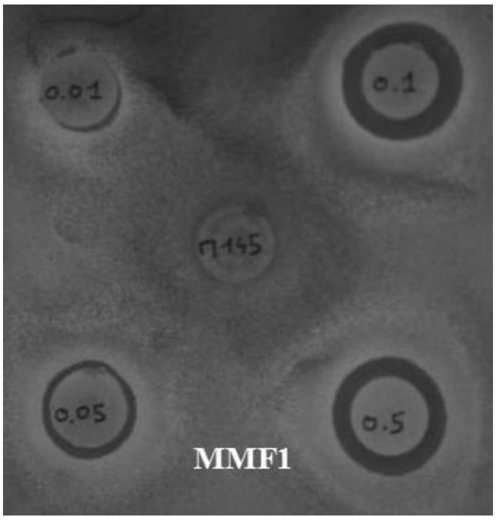

**Supplementary Fig. 3 | *In vivo* assay for induction of methylenomycin production upon addition of increasing amounts of MMF1 signalling molecules to growing mycelia of the MMF non-producing strain *S. coelicolor* W81.** Methylenomycin production was detected by growth inhibition of the methylenomycin-sensitive strain *S. coelicolor* M145.

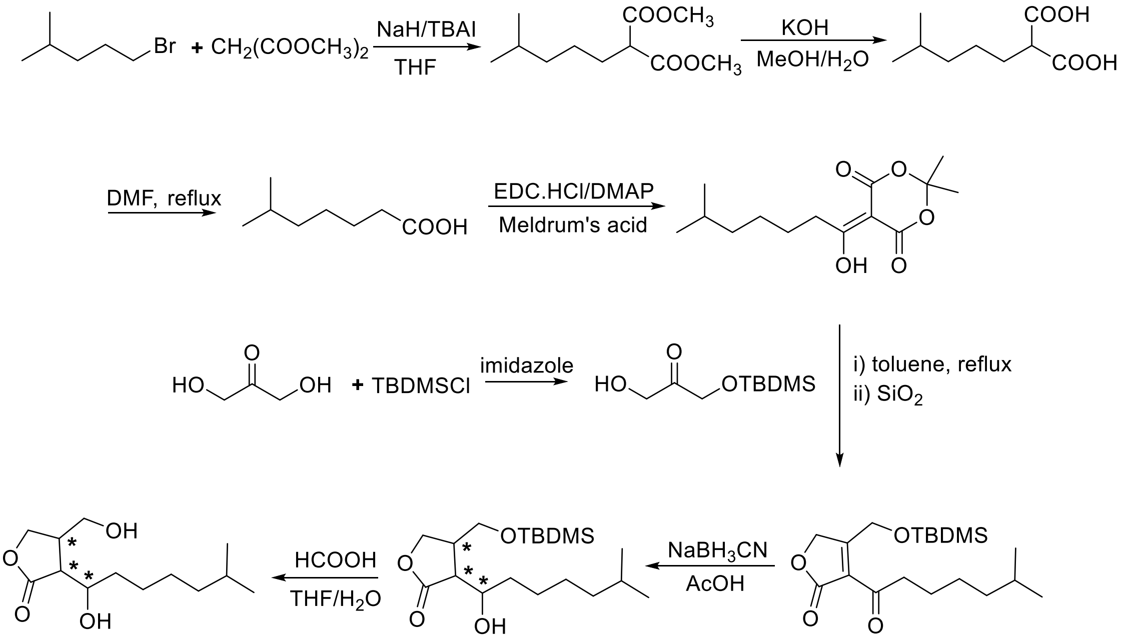

**Supplementary Fig. 4 | Synthetic route to SCB1.**

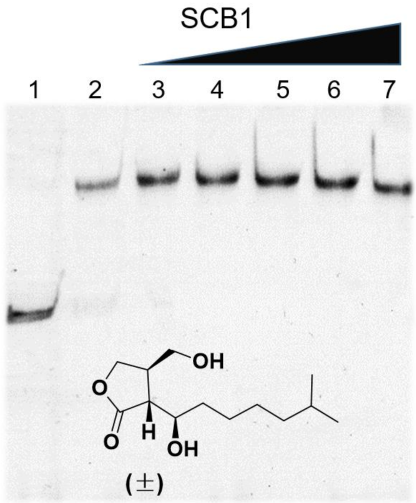

**Supplementary Fig. 5 | Interaction of MmfR with the DNA fragments corresponding to the *mmfL-mmfR* intergenic region (194 bp) in response to increasing amounts of SCB1.** Lane 1: isolated DNA fragments (0.1 pmol); lane 2: DNA fragments mixed with MmfR (0.1 pmol and 1.8 pmol respectively); lanes 3 to 7: addition of increasing quantities of racemic SCB1 (0.8, 8, 20, 40 and 400 nmol respectively) to the DNA/protein complexes.

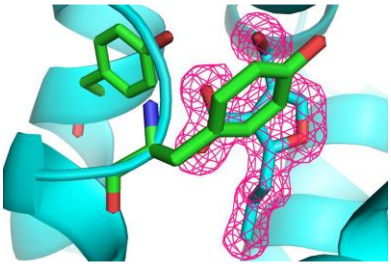

**Supplementary Fig. 6 | Close-up of the ligand binding site of the MmfR-AHFCA protein-ligand complex.** The SIGMAA-weighted 2mFo-DFc electron density surrounding the bound AHFCA ligand is shown in mesh representation, and two key conserved tyrosine residues (Y85 on the left, Y144 above the ligand) are shown in stick representation. The electron density map is contoured at the 1.5, which represents the RMS electron density for the unit cell. Contours further than 1.4 Å away from any of the displayed atoms have been removed for clarity. The figure was drawn with PyMOL.

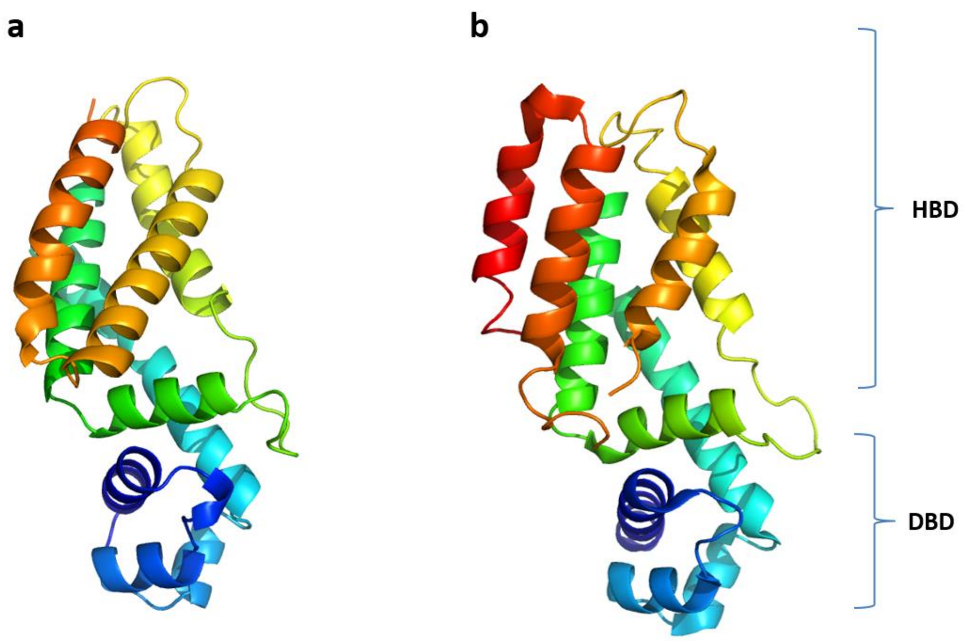

**Supplementary Fig. 7 | Cartoon representations of MmfR and CprB monomers. a**, MmfR. **b**, CprB. Colour ramped from blue to red from the N- to C- terminus. DBD: DNA-binding domain; HBD: hormone-binding domain.

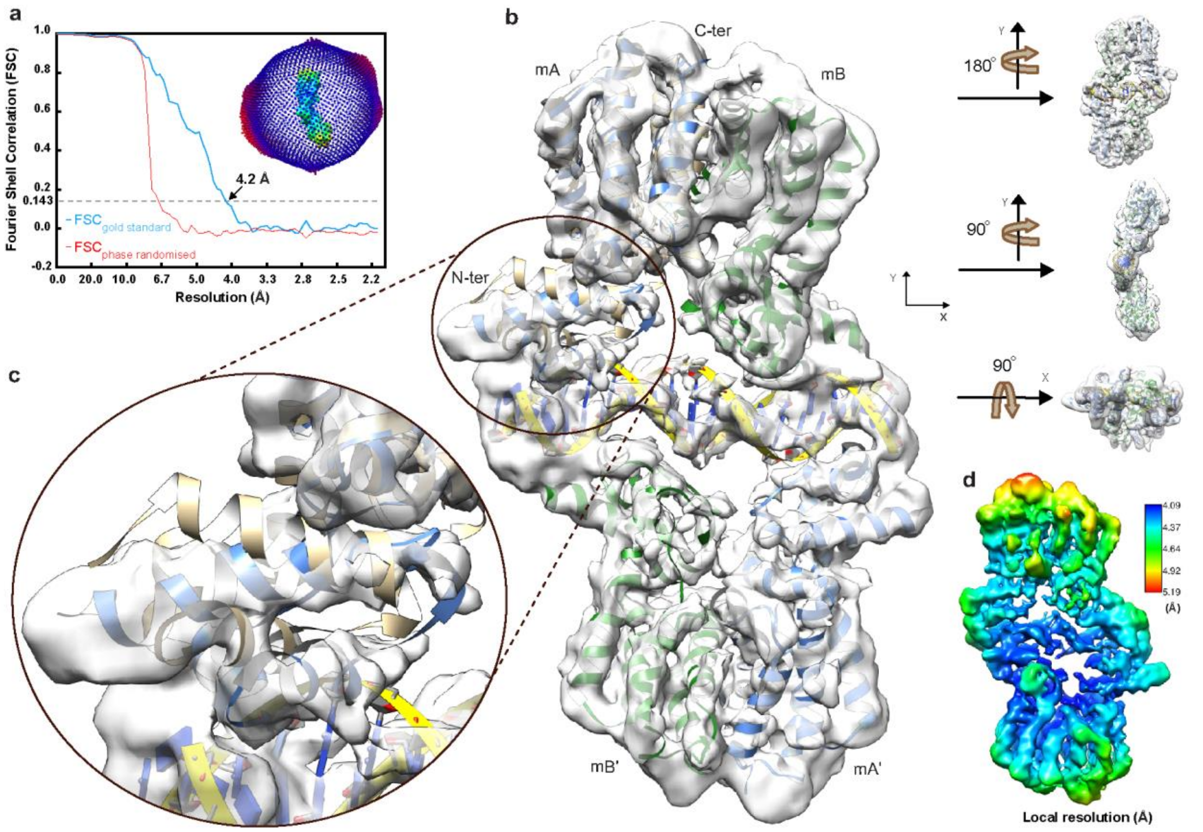

**Supplementary Fig. 8 | Data quality, overall view of model and cryo-EM map fit for the MmfR-MARE1 complex**. **a**, Relion corrected Fourier shell correlation (FSC) curve of protein-DNA complex map. The inset shows the angular distribution of the particle projections. The length of the projection is a direct measure of the number of assigned particles in each direction. **b**, Model construction by fitting the coordinates for the MmfR-MMF2 complex (pink and blue cartoon) into the cryo-EM density map. Different views of the cryo-EM density maps with the protein-DNA complex modelled into it are shown to the right. **c**, Zoomed-in view of the DBD showing how differently it is oriented in the MmfR-MM2 complex compared to the MmfR-MARE1 complex. **d**, Local resolution values projected onto the experimental density map.

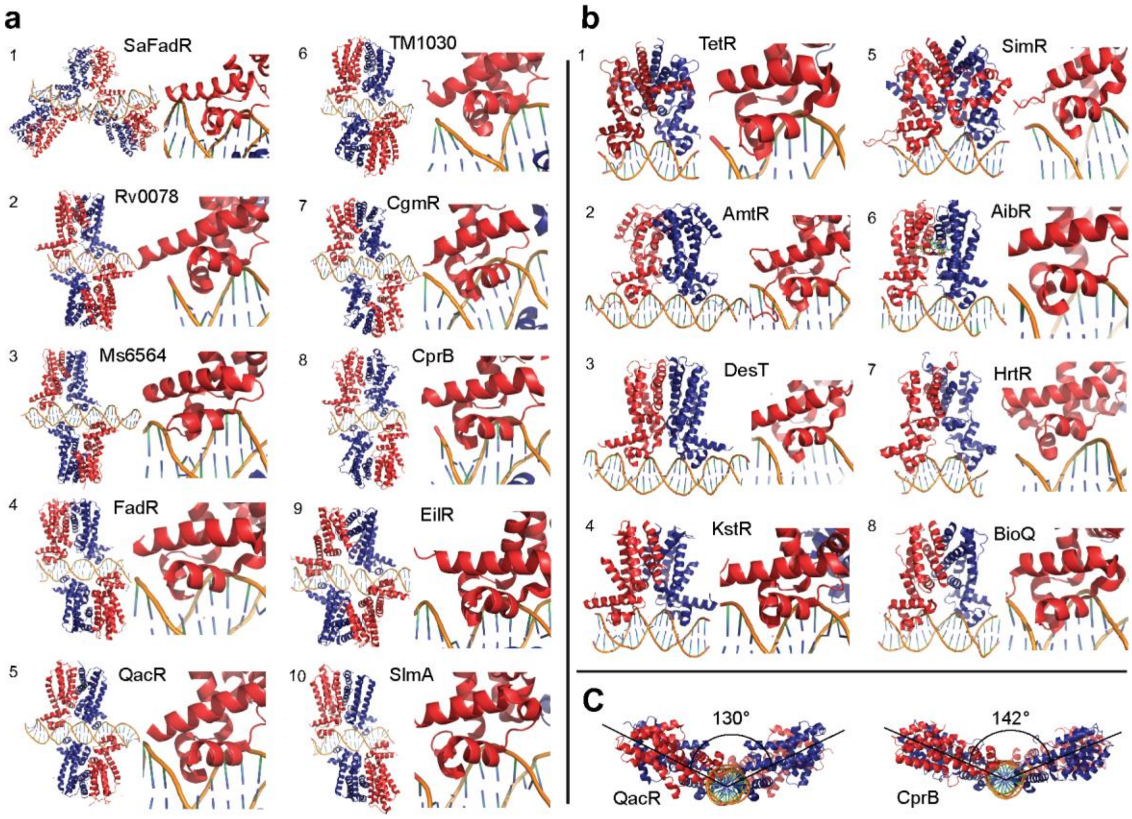

**Supplementary Fig. 9 | X-ray crystal structures of TFTRs in complex with their operators. a**, Dimer-of-dimer DNA binding and **b**, dimer DNA binding TFTRs. PDB entries for **a** (1 – 10) 6EN8, 6C31, 4JL3, 5GPC, 1JT0, 4I6Z, 2YVH, 4PXI, 5VL9 and 4GCT and, **b** (1 – 8) are 1QPI, 5DY0, 3LSP, 5UA1, 3ZQL, 5K7Z, 3VOK and 5YEJ respectively. **c**, Emphasizing the angle between the planes dissecting monomers bound to opposite sides of DNA in representative members of dimer-of-dimer DNA binding TFTRs.

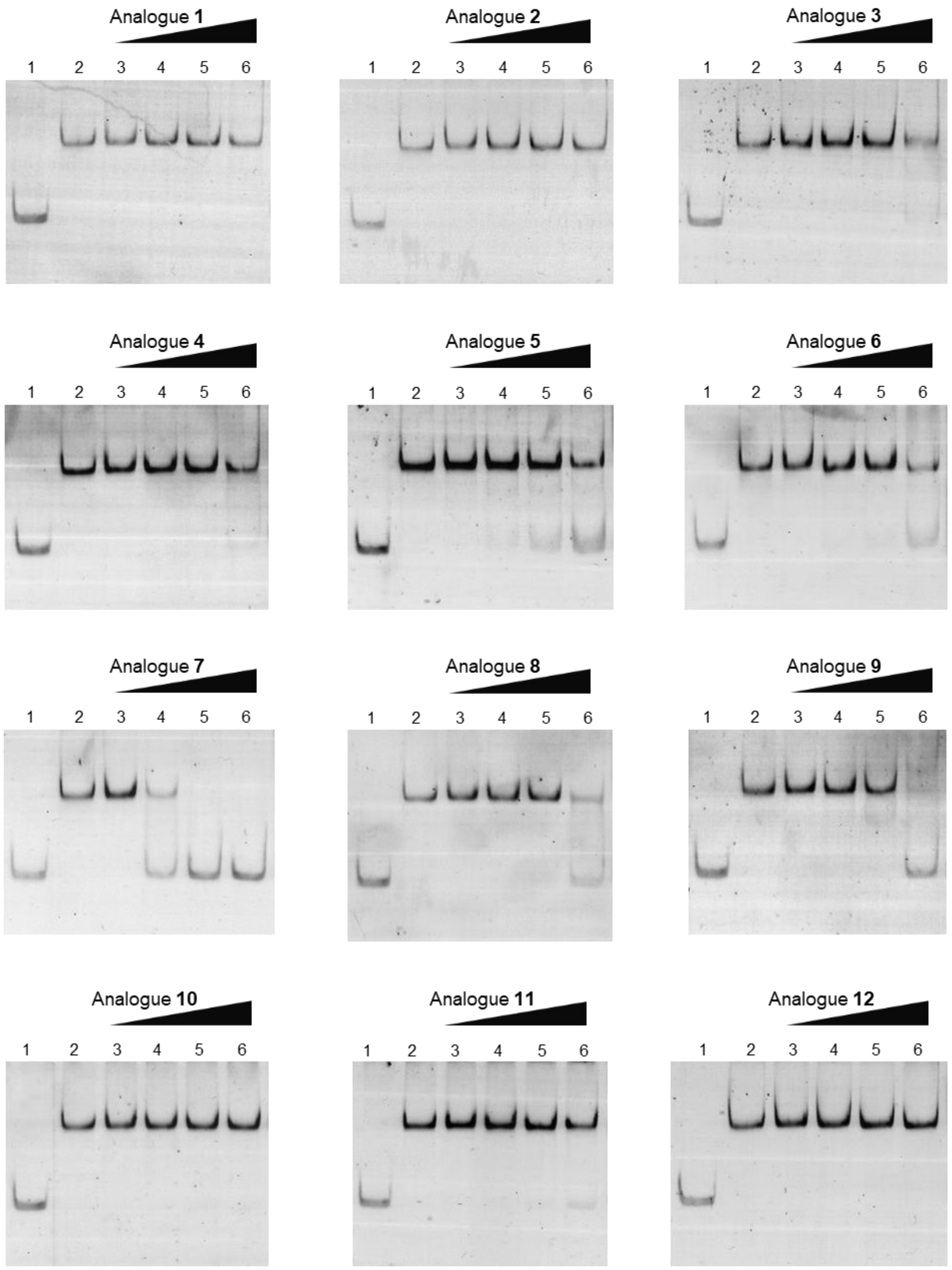

**Supplementary Fig. 10 | Interaction of MmfR with the DNA fragments corresponding to the *mmfL-mmfR* intergenic region (194 bp) in response to increasing amounts of synthetic MMF analogues.** Lane 1: isolated DNA fragments (0.1 pmol); lane 2: DNA fragments mixed with MmfR (0.1 pmol and 1.8 pmol respectively); lanes 3 to 6: addition of increasing quantities of MMF analogues (0.8, 8, 20 and 100 nmol respectively) to the DNA/protein complexes.

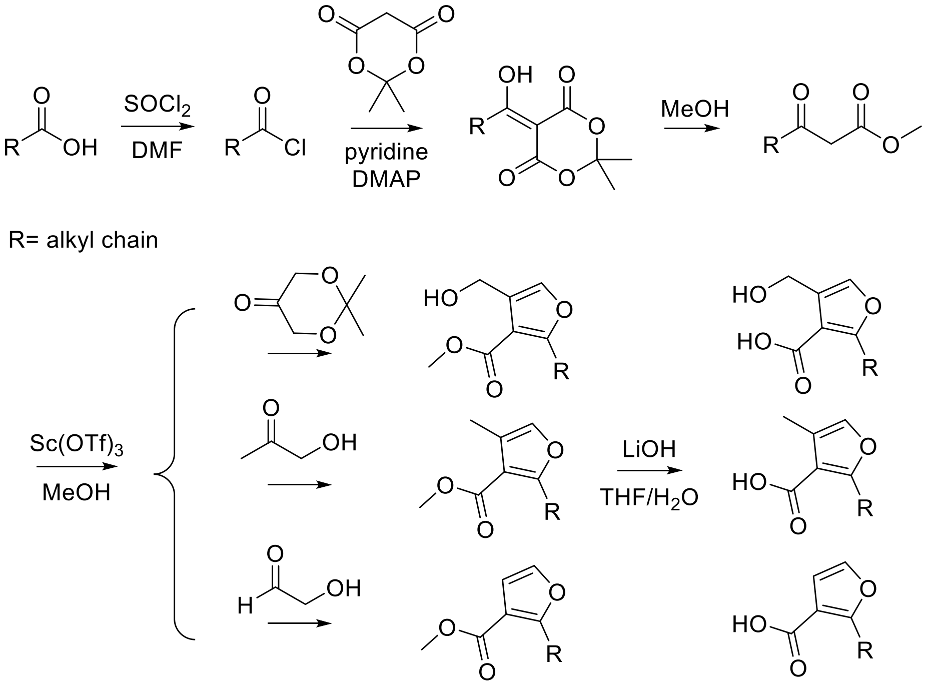

**Supplementary Fig. 11 | General synthetic route to MMFs and most analogues.**

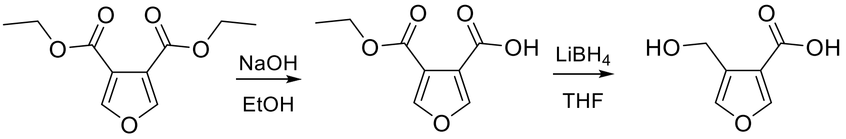

**Supplementary Fig. 12 | Synthetic route for MMF analogue 1 lacking a 2-alkyl group.**

**Supplementary Table 1** Comparison of the ability of the five naturally occurring MMFs to induce production of methylenomycin A in *S. coelicolor* W81.

| Compound | Structure | R= | Amount (μg) | | | | |
|---|---|---|---|---|---|---|---|
| | | | 0.05 | 0.1 | 0.5 | 5 | 50 |
| MMF1 | | 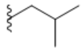 | o | o | o | o | o |
| MMF2 | | 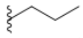 | x | x | o | o | o |
| MMF3 | 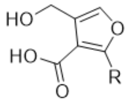 | 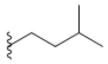 | x | x | o | o | o |
| MMF4 | | 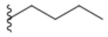 | x | o | o | o | o |
| MMF5 | | 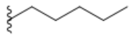 | x | o | o | o | o |

Key: o methylenomycin production was observed; x no methylenomycin production.

**Supplementary Table 2** Crystallographic data collection, phasing and refinement statistics.

| | **Native (PDB** 6SRM**)** | **Liganded (PDB** 6SRN**)** | **Se-Met derivative** |
|---|---|---|---|
| **Data collection** | | | |
| Beamline | Diamond I24 | Diamond IO4 | Diamond IO3 |
| Detector | Pilatus 6M | ADSC | Pilatus 6M-F |
| Wavelenth (Å) | 0.9686 | 0.9795 | 0.9793 |
| Space group | C222 | $P4_12_12$ | $P4_12_12$ |
| Unit cell a,b,c (Å) | 67.50, 115.72, 53.09 | 67.21, 67.21, 92.82 | 67.04, 67.04, 93.91 |
| Resolution (Å) | 53.61-1.5 (1.58-1.5) | 42.3-1.5 (1.58-1.5) | 93.9-2.2 (2.32-2.2) |
| Observations | 129,867 (17,251) | 408,083 (58,085) | 145,517 (20,830) |
| Unique reflections | 29,985 (1,256) | 34,705 (1,507) | 11,459 (1,623) |
| $I/\sigma(I)$ | 16.0 (2.2) | 17.9 (2.2) | 18.5 (4.5) |
| $R_{sym}$ | 0.053 (0.659) | 0.082 (1.189) | 0.137 (0.768) |
| Completeness (%) | 99.3 (99.6) | 99.9 (99.6) | 100 (100) |
| **Phasing (SAD)** | | | |
| Mean figure of merit | | | 0.30 |
| Sites found | | | 7 |
| **Refinement** | | | |
| Non-hydrogen atoms | | | |
| Protein | 1,497 | 1,480 | |
| Solvent (water/glycerol) | 189 | 256/6 | |
| Ligand (AHFCA) | | 13 | |
| $R_{cryst}$ | 0.214 (0.352) | 0.174 (0.267) | |
| Reflections used | 28,803 (1,973) | 33,288 (2,379) | |
| $R_{free}$ | 0.243 (0.355) | 0.207 (0.240) | |
| Reflections used | 1,182 (74) | 1,417 (90) | |
| $R_{cryst}$ (all data) | 0.215 | 0.175 | |
| Mean B value (Å$^2$) | 16.7 | 15.2 | |
| Rmsds from ideal values | | | |
| Bonds (Å) | 0.015 | 0.014 | |
| Angles (°) | 1.75 | 1.9 | |
| DPI coordinate error (Å) | 0.090 | 0.068 | |

Numbers in parentheses refer to values in the highest resolution shell.
$R_{sym} = \Sigma_j\Sigma_h|I_{h,j} - <I_h>|/\Sigma_j\Sigma_h<I_h>$ where $I_{h,j}$ is the jth observation of reflection h, and $<I_h>$ is the mean intensity of that reflection.
$R_{cryst} = \Sigma||F_{obs}|-|F_{calc}||/\Sigma|F_{obs}|$ where $F_{obs}$ and $F_{calc}$ are the observed and calculated structure factor amplitudes, respectively.
$R_{free}$ is equivalent to $R_{cryst}$ for a 4% subset of reflections not used in the refinement.
DPI refers to the diffraction component precision index.

**Supplementary Table 3** Cryo-EM data collection and processing for the MmfR-MARE1 complex.

| | (Data set 1) | (Data set 2) |
|---|---|---|
| **Data collection and processing** | | |
| Magnification | 165,000 | 165,000 |
| Voltage (kV) | 300 | 300 |
| Electron exposure (e–/Å$^2$) | 50 | 50 |
| Defocus range (μm) | 0.5 | 0.5 |
| Pixel size (Å) | 0.84 | 0.84 |
| Initial particle images (no.) | 181,409 | 292,512 |
| Final particle images (no.) | 311088 | |
| Symmetry imposed | C1 | |
| Map resolution (Å) | 4.5 | |
| FSC threshold | 0.143 | |
| Map sharpening *B* factor (Å$^2$) | -94.02 | |
| Initial model used (PDB code) | 4PXI | |
| Model composition | | |
| Non-hydrogen atoms | 6777 | |
| Protein residues | 756 | |
| DNA residues | 38 | |
| Ramachandran plot | | |
| Favored (%) | 97.86 | |
| Allowed (%) | 2.14 | |
| Disallowed (%) | 0.0 | |
| EMDB | EMD-20781 | |

**Supplementary Table 4** Comparison of the ability of the synthesized MMF analogues to induce production of methylenomycin A in *S. coelicolor* W81.

| Compound | Structure | R= | Amount (μg) | | | | |
|---|---|---|---|---|---|---|---|
| | | | 0.05 | 0.1 | 0.5 | 5 | 50 |
| MMF1 | | 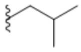 | o | o | o | o | o |
| **1** | | 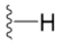 | x | x | x | x | x |
| **2** | | 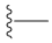 | x | x | x | x | x |
| **3** | | 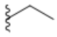 | x | x | x | o | o |
| **4** | 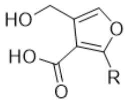 | 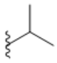 | x | x | o | o | o |
| **5** | | 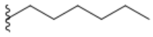 | x | x | o | o | o |
| **6** | | 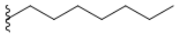 | x | x | x | o | o |
| **7** | | 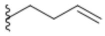 | x | o | o | o | o |
| **8** | | 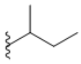 | x | x | o | o | o |
| **9** | | 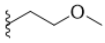 | x | x | x | x | o |
| **10** | 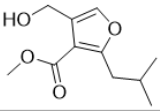 | | x | x | x | x | x |
| **11** | 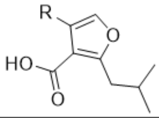 | 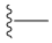 | x | x | o | o | o |
| **12** | | 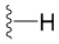 | x | x | x | o | ND |
| SCB1 | 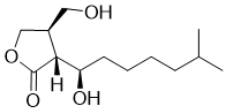 | | x | x | x | x | x |

Key: o methylenomycin production was observed; x no methylenomycin production; ND not determined.

**Supplementary Table 5** Strains and plasmids used in this study.

| Strains / Plasmids | Relevant properties | Source or Reference |
|---|---|---|
| **Strains** | | |
| ***Streptomyces coelicolor*** | | |
| M145 | $SCP1^-$, $SCP2^-$ (Methylenomycin-sensitive) | [1] |
| W81 | M145 with pCC002::*attB* (MMF minus, methylenomycin minus) | [2] |
| ***E. coli*** | | |
| TOP10 | One shot chemically competent cells | Invitrogen |
| BL21 Star (DE3) | $F^-$*ompT hsdS*$_B$ ($r_B^-$, $m_B^-$) *galdcmrne*131 (DE3) T7 promoter based expression system | Invitrogen |
| B834 (DE3) | Met auxotrophic strain for production of Se-Met-MmfR | Novagen |
| **Plasmids** | | |
| pET151/D-TOPO | T7 expression vector, ampicillin resistance | Invitrogen |
| pET151-*mmfR* | - | This study |
| pET151-*mmfR* Q130E | - | This study |
| pET151-*mmfR* Y85F | - | This study |
| pET151-*mmfR* Y144F | - | This study |
| C73_787 | Integrative cosmid containing the full methylenomycin cluster including the MMFs biosynthetic genes | [3] |
| pCC002 | C73_787 with *mmfL→P*::*apr* | [2] |

**Supplementary Table 6** Primers used for cloning of *mmfR*, mutagenesis and PCR amplification of the intergenic regions for EMSAs and oligonucleotides for hairpin DNA formation. Fw: forward primer, Rev: reverse primer.

| Target | Sequences (5'-3') | Details |
|---|---|---|
| ***Protein overproduction*** | | |
| *mmfR* | Fw: CACCATGACGAGCGCCCAACAAC<br>Rev: TCAGGCGCGGAGAGCGAAG | Cloning of *mmfR* gene (SCP1.242c – 642 bp) into pET151/D-TOPO vector |
| ***Site directed mutagenesis*** | | |
| *mmfR*<br>(Q130E) | Fw: TGCCCGGCTGGAGAGTGAGCG<br>Rev: CCGGCCTGCATCACGGGG | Q5; New England Biolabs |
| *mmfR*<br>(Y85F) | Fw: GAGGAGCACTTCGCGCGCTGG<br>Rev: CACCACGGCGATGGCCAG | Q5; New England Biolabs |
| *mmfR*<br>(Y144F) | Fw: CCCCTGCCCTTCGTGGACTGG<br>Rev: CAGCTCCGCGTCGATGAAG | Q5; New England Biolabs |
| ***PCR amplification of intergenic regions*** | | |
| *mmfL-mmfR*<br>194 bp | Fw: GGCTGCCTTCCTTCGTGTG<br>Rev: AGGGGCGCTACATCTCCCG | Entire intergenic region between *mmfL* and *mmfR* |
| *mmyB-mmyY*<br>230 bp | Fw: GGTGAACTCCTTCGGCGAG<br>Rev: GGCGCCTCACAGTGTCAAAC | Entire intergenic region between *mmyB* and *mmyY* |
| *mmfL-mmfR*<br>100 bp | Fw: CACGGAAACCCATTGCATAATACC<br>Rev: GGACCACCGGCTGGCTTGC | Partial intergenic sequence between *mmfL* and *mmfR* |
| *mmyB-mmyY*<br>98 bp | Fw: TCCAAACACCGAGGCCCCG<br>Rev: GGCGCCTCACAGTGTCAAAC | Partial intergenic sequence between *mmyB* and *mmyY* |
| ***Hairpin DNA formation*** | | |
| MARE1<br>(*mmfL-mmfR*) | GCAATATACCTGCGGGAAGGTATTATGCGAG<br>GCATAATACCTTCCCGCAGGTATATTGC | For hairpin DNA containing the operator between *mmfL* and *mmfR* |
| MARE2<br>(*mmyB-mmyY*) | GCAAAAAACCTTCGGGAAGGTTTGACGCGAG<br>GCGTCAAACCTTCCCGAAGGTTTTTTGC | For hairpin DNA containing the operator between *mmyB* and *mmyY* |

### Chemical synthesis

#### *General experimental details*

Anhydrous toluene, tetrahydrofuran (THF), N,N-dimethylformamide (DMF) and dichloromethane (DCM) were obtained by distillation from calcium hydride under argon and stored over activated 4 Å molecular sieves. All other reagents and solvents were used as supplied (Sigma-Aldrich, VWR or Fisher Scientific). Flash column chromatography was conducted on Sigma-Aldrich silica gel (40-63 µm, 60 Å). Thin layer chromatography (TLC) was carried out on aluminium backed sheets pre-coated with Merck silica gel 60 $F_{254}$ and visualised by UV radiation, phosphomolybdic acid or potassium permanganate. The $^{1}$H- and $^{13}$C-NMR spectra were obtained using a Bruker DPX 300 or DPX 400 spectrometer. Chemical shifts ($\delta$) are given in ppm with reference to the residual solvent peak. Coupling constants ($J$) are rounded to the nearest 0.5 Hertz (Hz). Multiplicities are given as multiplet (m), singlet (s), doublet (d), triplet (t), quartet (q), quintet (quin), sextet (sext), septet (sept) and nonet (non). Low resolution ESI mass spectra were recorded using an Agilent 6130B single quadrupole spectrometer. High resolution mass spectra (HRMS) were recorded on a Bruker MaXis impact mass spectrometer.

#### *Synthesis of MMFs and analogues*

Most of the MMFs and analogues with alteration to the alkyl chains were synthesized according to literature procedures [4, 5]. The general synthetic route is shown in **Supplementary Figure 11**. It includes preparation of the desired β-ketoester [4], and subsequent scandium triflate-catalysed condensation with the desired ketone prepared according to published literatures: dihydroxyacetone acetonide [6], hydroxylacetone, hydroxyacetaldehyde, 1-amino-3-hydroxypropan-2-one [7] or mercaptopropanone [8].

MMF analogue **10** is the methyl ester intermediate during MMF1 synthesis. The MMF analogue **1** without an alkyl chain was produced using a reported method of mono-hydrolysis of the di-ester starting material by dilute reaction condition [9] followed by selective reduction of the ester group [10] in **Supplementary Figure 12**.

#### *General procedure for synthesis of β-ketoesters*

Thionyl chloride (2 eq.) and a catalytic amount of DMF were slowly added to starting acid (1 eq.) under argon. The mixture was stirred overnight at room temperature. The volatile material was removed *in vacuo* using a sodium hydroxide trap. The resulting crude material was used without further purification. To a 0.5 M solution of Meldrum's acid (1 eq.) in dry DCM under argon, dry pyridine (2 eq.) and 4-dimethylaminopyridine (0.2 eq.) were added and the mixture was stirred for 10 min at room temperature before being cooled to 0 °C. Then 1 eq. of the desired acid chloride (or the previous crude mixture) was added dropwise and the reaction was warmed to room temperature and stirred overnight. The resulting mixture was washed with 1 M HCl and water. The organic layer was dried over $MgSO_4$, filtered and concentrated *in vacuo*. The resulting dark-yellow oil was diluted in methanol to a final concentration of 0.5 M. The reaction was refluxed for 5 h before the solvent was removed *in vacuo*. The oil residue was purified by flash chromatography (pentane/diethyl ether = 6:1 *v/v*) to afford the desired β-ketoester as a pale yellow oil.

The β-ketoesters for **MMF4** and MMF analogue **2, 3** and **9** are commercially available and used as received.

**Methyl 5-methyl-3-oxohexanoate**

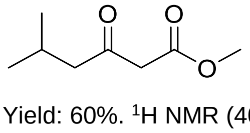

Yield: 60%. $^{1}$H NMR (400 MHz, $CDCl_3$) $\delta$: 0.83 (d, $J$ = 6.5 Hz, 6H), 2.05 (m, 1H), 2.32 (d, $J$ = 7.0 Hz, 2H), 3.35 (s, 2H), 3.63 (s, 3H). $^{13}$C NMR (100 MHz, $CDCl_3$) $\delta$: 202.3, 167.6, 52.1, 51.7, 49.3, 24.2, 22.3. HR-MS: *m/z* calculated for $C_8H_{14}O_3$ [M+Na]$^+$: 181.0835; found: 181.0833.

**Methyl 3-oxohexanoate**

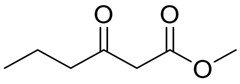

Yield: 55%. $^{1}H$ NMR (400 MHz, $CDCl_3$) $\delta$: 0.92 (t, $J$ = 7.5 Hz, 3H), 1.61 (sext, $J$ = 7.5 Hz, 2H), 2.54 (t, $J$ = 7.5 Hz, 2H), 3.47 (s, 2H), 3.71 (s, 3H). $^{13}C$ NMR (100 MHz, $CDCl_3$) $\delta$: 202.5, 167.5, 51.8, 48.6, 44.4, 16.6, 13.2. HR-MS: *m/z* calculated for $C_7H_{12}O_3$ $[M+Na]^+$: 167.0679; found: 167.0683.

**Methyl 6-methyl-3-oxoheptanoate**

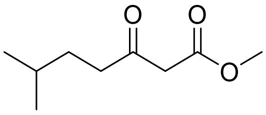

Yield: 21%. $^{1}H$ NMR (400 MHz, $CDCl_3$) $\delta$: 0.83 (d, $J$ = 6.0 Hz, 6H), 1.54 (m, 3H), 2.54 (m, 2H), 3.48 (s, 2H), 3.73 (s, 3H). $^{13}C$ NMR (100 MHz, $CDCl_3$) $\delta$: 202.9, 179.9, 167.8, 40.9, 33.4, 32.0, 27.4, 22.1. HR-MS: *m/z* calculated for $C_9H_{16}O_3$ $[M+Na]^+$: 195.0992; found: 195.0991.

**Methyl 3-oxooctanoate**

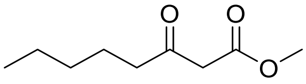

Yield: 80%. $^{1}H$ NMR (400 MHz, $CDCl_3$) $\delta$: 0.89 (t, $J$ = 7.5 Hz, 3H), 1.29 (m, 4H), 1.57 (br quin, $J$ = 7.5 Hz, 2H), 2.54 (t, $J$ = 7.5 Hz, 2H), 3.46 (s, 2H), 3.69 (s, 3H). $^{13}C$ NMR (100 MHz, $CDCl_3$) $\delta$: 202.3, 167.4, 51.5, 48.4, 42.3, 30.8, 22.7, 22.1, 13.4. HR-MS: *m/z* calculated for $C_9H_{16}O_3$ $[M+Na]^+$: 195.0997; found: 195.0995.

**Methyl 4-methyl-3-oxopentanoate**

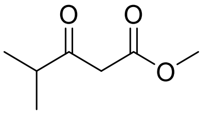

Yield: 38%. $^{1}H$ NMR (400 MHz, $CDCl_3$) $\delta$: 1.04 (d, $J$ = 7.0 Hz, 6H), 2.65 (sept, $J$ = 7.0 Hz, 1H), 3.45 (s, 2H), 3.63 (s, 3H). $^{13}C$ NMR (100 MHz, $CDCl_3$) $\delta$: 206.3, 197.7, 51.9, 46.7, 40.9, 17.6. HR-MS: *m/z* calculated for $C_7H_{12}O_3$ $[M+Na]^+$: 167.0679; found: 167.0979.

**Methyl 3-oxononanoate**

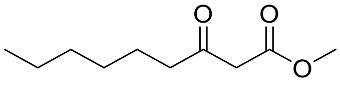

Yield: 30%. $^{1}H$ NMR (400 MHz, $CDCl_3$) $\delta$: 0.88 (m, 3H), 1.29 (m, 6H), 1.61 (m, 2H), 2.54 (t, $J$ = 7.5 Hz, 2H), 3.46 (s, 2H), 3.72 (s, 3H). $^{13}C$ NMR (100 MHz, $CDCl_3$) $\delta$: 202.8, 167.7, 52.03, 48.7, 42.8, 31.4, 28.5, 23.3, 22.3, 13.8. HR-MS: *m/z* calculated for $C_{10}H_{18}O_3$ $[M+Na]^+$: 209.1148; found: 209.1151.

**Methyl 3-oxodecanoate**

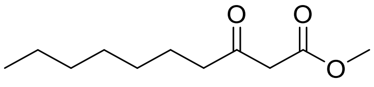

Yield: 85%. $^1$H NMR (400 MHz, $CDCl_3$) *δ*: 0.81 (t, *J* = 7.0 Hz, 3H), 1.22 (m, 8H), 1.53 (m, 2H), 2.47 (t, *J* = 7.5 Hz, 2H), 3.39 (s, 2H), 3.67 (s, 3H). $^{13}$C NMR (100 MHz, $CDCl_3$) δ: 202.9, 167.7, 52.2, 49.0, 43.0, 31.6, 29.0, 29.0, 23.5, 22.6, 14.0. HR-MS: *m/z* calculated for $C_{11}H_{20}O_3$ $[M+Na]^+$: 223.1305; found: 223.1304.

**Methyl 3-oxohept-6-enoate**

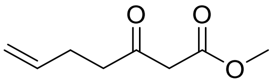

Yield: 51%. $^1$H NMR (400 MHz, $CDCl_3$) *δ*: 2.17 (m, 2H), 2.50 (m, 2H), 3.31 (m, 2H), 3.56 (m, 3H), 4.86 (m, 2H), 5.64 (m, 1H). $^{13}$C NMR (100 MHz, $CDCl_3$) *δ*: 201.7, 167.4, 136.5, 115.3, 51.9, 48.7, 41.7, 27.2. HR-MS: *m/z* calculated for $C_8H_{12}O_3$ $[M+Na]^+$: 179.0696; found: 179.0696.

**Methyl 4-methyl-3-oxohexanoate**

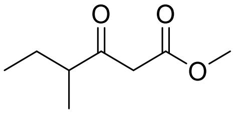

Yield: 34%. $^1$H NMR (400 MHz, $CDCl_3$) *δ*: 0.73 (m, 3H), 0.94 (m, 3H), 1.26 (m, 1H), 1.54 (m, 1H), 2.42 (m, 1H), 3.35 (s, 2H), 3.55 (s, 3H). $^{13}$C NMR (100 MHz, $CDCl_3$) *δ*: 206.2, 167.7, 51.9, 47.8, 47.2, 25.3, 15.1, 11.1. HR-MS: *m/z* calculated for $C_8H_{14}O_3$ $[M+Na]^+$: 181.0843; found: 181.0843.

***General procedure for furan cyclisation***

The desired ketone (1 eq.) and β-ketoester (1 eq.) were dissolved in methanol to obtain a 0.5 M solution. Scandium III triflate (0.1 eq.) was added and allowed to stir overnight at room temperature. The solvent was removed *in vacuo* and the residue was purified by flash chromatography (pentane/diethyl ether = 4:1 *v/v*) giving the desired furan as a pale yellow oil.

**Methyl 4-(hydroxymethyl)-2-isobutylfuran-3-carboxylate (MMF analogue 10)**

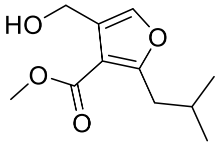

Yield: 76%. $^1$H NMR (400 MHz, $CDCl_3$) *δ*: 0.91 (d, *J* = 6.5 Hz, 6H), 2.04 (m, 1H), 2.81 (d, *J* = 7.0 Hz, 2H), 3.82 (s, 3H), 4.20 (br s, 1H), 4.58 (s, 2H), 7.27 (s, 1H). $^{13}$C NMR (100 MHz, $CDCl_3$) *δ*: 165.1, 163.6, 138.3, 126.0, 112.2, 55.8, 51.1, 36.5, 28.1, 22.1. HR-MS: *m/z* calculated for $C_{11}H_{16}O_4$ $[M+Na]^+$: 235.0941; found: 235.0941.

**Methyl 4-(hydroxymethyl)-2-propylfuran-3-carboxylate**

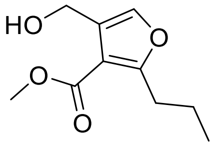

Yield: 84%. $^1$H NMR (400 MHz, $CDCl_3$) *δ*: 0.94 (t, *J* = 7.5 Hz, 3H), 1.68 (sext, *J* = 7.5 Hz, 2H), 2.91 (t, *J* = 7.5 Hz, 2H), 3.85 (s, 3H), 3.91 (br s, 1H), 4.56 (s, 2H), 7.25 (s, 1H). $^{13}$C NMR (100 MHz, $CDCl_3$) *δ*: 165.4, 164.5, 138.3, 125.9, 111.9, 55.8, 51.4, 29.9, 21.2, 13.6. HR-MS: *m/z* calculated for $C_{10}H_{14}O_4$ $[M+Na]^+$: 221.0784; found: 221.0786.

**Methyl 4-(hydroxymethyl)-2-isopentylfuran-3-carboxylate**

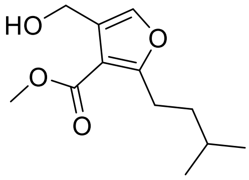

Yield: 83%. $^1$H NMR (400 MHz, $CDCl_3$) *δ:* 0.92 (d, *J* = 6.0 Hz, 6H), 1.54 (m, 3H), 2.81 (t, *J* = 7.5 Hz, 2H), 4.13 (br s, 1H), 3.82 (s, 3H), 4.56 (s, 2H), 7.24 (s, 1H). $^{13}$C NMR (100 MHz, $CDCl_3$) *δ*: 165.1, 164.6, 138.1, 126.1, 111.4, 55.8, 51.1, 36.7, 27.5, 25.9, 23.0. HR-MS: *m/z* calculated for $C_{12}H_{18}O_4$ $[M+Na]^+$: 249.1097; found: 249.1093.

**Methyl 2-butyl-4-(hydroxymethyl)furan-3-carboxylate**

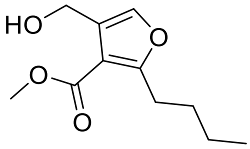

Yield: 80%. $^1$H NMR (400 MHz, $CDCl_3$) *δ*: 0.92 (t, *J* = 7.5 Hz, 3H), 1.35 (sext, *J* = 7.5 Hz, 2H), 1.63 (quin, *J* = 7.5 Hz, 2H), 2.94 (t, *J* = 7.5 Hz, 2H), 3.85 (s, 3H), 3.91 (br s, 1H), 4.56 (s, 2H), 7.24 (s, 1H). $^{13}$C NMR (100 MHz, $CDCl_3$) *δ*: 165.4, 164.7, 138.2, 125.9, 111.8, 55.8, 51.4, 29.9, 27.7, 22.2, 13.6. HR-MS: *m/z* calculated for $C_{11}H_{16}O_4$ $[M+Na]^+$: 235.0941; found: 235.0940.

**Methyl 4-(hydroxymethyl)-2-pentylfuran-3-carboxylate**

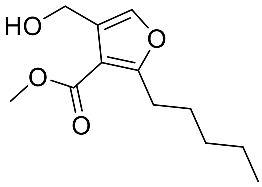

Yield: 76%. $^1$H NMR (400 MHz, $CDCl_3$) *δ*: 0.89 (t, *J* = 7.0 Hz, 3H), 1.35 (m, 4H), 1.66 (q, *J* = 7.5 Hz, 2H), 2.93 (t, *J* = 7.5 Hz, 2H), 3.85 (s, 3H), 3.87 (br s, 1H), 4.56 (s, 2H), 7.24 (s, 1H). $^{13}$C NMR (100 MHz, $CDCl_3$) *δ*: 165.4, 164.7, 138.2, 125.9, 111.8, 55.8, 51.4, 31.2, 28.0, 27.5, 22.2, 13.8. HR-MS: *m/z* calculated for $C_{12}H_{18}O_4$ $[M+Na]^+$: 249.1097; found: 249.1097.

***tert*-Butyl 4-(hydroxymethyl)-2-methylfuran-3-carboxylate**

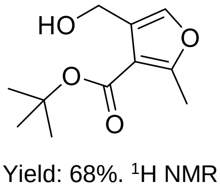

Yield: 68%. $^1$H NMR (400 MHz, $CDCl_3$) *δ*: 1.47 (s, 9H), 2.40 (s, 3H), 4.01 (br s, 1H), 4.43 (s, 2H), 7.09 (s, 1H). $^{13}$C NMR (100 MHz, $CDCl_3$) *δ*: 164.6, 160.0, 137.7, 126.1, 113.9, 81.5, 55.8, 28.2, 14.3. HR-MS: *m/z* calculated for $C_{11}H_{16}O_4$ $[M+Na]^+$: 235.0941; found: 235.0940.

**Methyl 2-ethyl-4-(hydroxymethyl)furan-3-carboxylate**

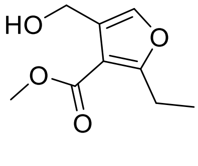

Yield: 85%. $^1$H NMR (400 MHz, $CDCl_3$) *δ*: 1.22 (t, *J* = 7.5 Hz, 3H), 2.95 (q, *J* = 7.5 Hz, 2H), 3.84 (s, 3H), 4.15 (br s, 1H), 4.56 (s, 2H), 7.25 (s, 1H). $^{13}$C NMR (100 MHz, $CDCl_3$) *δ*: 165.3, 165.0, 138.2, 126.0, 111.1, 55.8, 51.3, 21.5, 11.9. HR-MS: *m/z* calculated for $C_9H_{12}O_4$ $[M+Na]^+$: 207.0628; found: 207.0624.

**Methyl 4-(hydroxymethyl)-2-isopropylfuran-3-carboxylate**

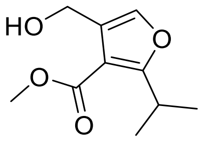

Yield: 79%. $^1$H NMR (400 MHz, $CDCl_3$) *δ*: 1.25 (d, *J* = 7.0 Hz, 6H), 3.69 (sept, *J* = 7.0 Hz, 1H), 3.77 (br s, 1H), 3.86 (s, 3H), 4.56 (s, 2H), 7.25 (s, 1H). $^{13}$C NMR (100 MHz, $CDCl_3$) *δ*: 168.5, 165.5, 138.2, 125.8, 110.4, 55.8, 51.6, 27.6, 20.6. HR-MS: *m/z* calculated for $C_{10}H_{14}O_4$ $[M+Na]^+$: 221.0784; found: 221.0787.

**Methyl 2-hexyl-4-(hydroxymethyl)furan-3-carboxylate**

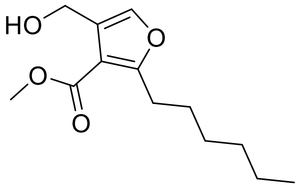

Yield: 40%. $^1$H NMR (400 MHz, $CDCl_3$) *δ*: 0.79 (m, 3H), 1.21 (m, 6H), 1.66 (m, 2H), 2.83 (t, *J* = 7.5 Hz, 2H), 3.76 (s, 3H), 3.86 (br s, 1H), 4.47 (s, 2H), 7.14 (s, 1H). $^{13}$C NMR (100 MHz, $CDCl_3$) *δ*: 165.4, 164.6, 138.2, 125.7, 111.8, 55.6, 51.5, 31.4, 28.4, 27.9, 22.5, 13.7. HR-MS: *m/z* calculated for $C_{13}H_{20}O_4$ $[M+Na]^+$: 263.1259; found: 263.1260.

**Methyl 2-heptyl-4-(hydroxymethyl)furan-3-carboxylate**

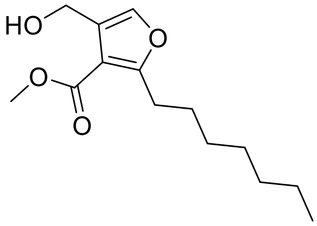

Yield: 70%. $^1$H NMR (400 MHz, $CDCl_3$) *δ*: 0.86 (t, *J* = 7.0 Hz, 3H), 1.27 (m, 8H), 1.62 (m, 2H), 2.90 (t, *J* = 7.5 Hz, 2H), 3.70 (br s, 1H), 3.85 (s, 3H), 4.53 (s, 2H), 7.22 (s, 1H). $^{13}$C NMR (100 MHz, $CDCl_3$) *δ:* 165.8, 165.0, 138.4, 129.9 112.1, 55.8, 51.8, 31.8, 29.2, 29.0, 28.3, 28.0, 22.7, 14.2. HR-MS: *m/z* calculated for $C_{14}H_{22}O_4$ $[M+Na]^+$: 277.1410; found: 277.1408.

**Methyl 2-(but-3-enyl)-4-(hydroxymethyl)furan-3-carboxylate**

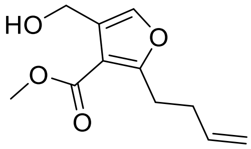

Yield: 82%. $^1$H NMR (400 MHz, $CDCl_3$) *δ*: 2.41 (q, *J* = 7.5 Hz, 2H), 3.04 (t, *J* = 7.5 Hz, 2H), 3.70 (br s, 1H), 3.87 (s, 3H), 4.56 (s, 2H), 4.98 (dd, *J* = 10.5 and 1.5 Hz, 1H), 5.04 (dd, *J* = 17.0 and 1.5 Hz, 1H), 5.82 (dddd, *J* = 17.0, 10.0, 6.5 and 6.5 Hz, 1H),7.25 (s, 1H). $^{13}$C NMR (100 MHz, $CDCl_3$) *δ*: 165.5, 163.7, 138.5, 136.9, 125.9, 115.5, 112.3, 55.8, 51.7, 31.9, 27.8. HR-MS: *m/z* calculated for $C_{11}H_{14}O_4$ $[M+Na]^+$: 233.0784; found: 233.0780.

**Methyl 2-sec-butyl-4-(hydroxymethyl)furan-3-carboxylate**

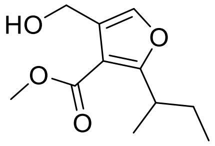

Yield: 68%. $^{1}H$ NMR (400 MHz, $CDCl_3$) *δ*: 0.72 (t, *J* = 7.5 Hz, 3H), 1.12 (d, *J* = 7.5 Hz, 3H), 1.55 (quin, *J* = 7.0 Hz, 2H), 3.41 (sext, *J* = 7.0 Hz, 1H), 3.74 (s, 3H), 3.87 (br s, 1H), 4.47 (s, 2H), 7.17 (s, 1H). $^{13}C$ NMR (100 MHz, $CDCl_3$) *δ*: 166.7, 164.6, 137.4, 124.9, 110.6, 55.0, 50.5, 33.3, 27.4, 17.5, 10.9. HR-MS: *m/z* calculated for $C_{11}H_{16}O_4$ $[M+Na]^+$: 235.0941; found: 235.0944.

**Methyl 4-(hydroxymethyl)-2-(2-methoxyethyl)furan-3-carboxylate**

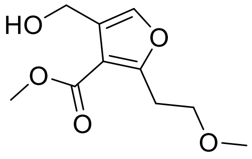

Yield: 75%. $^{1}H$ NMR (400 MHz, $CDCl_3$) *δ*: 3.22 (m, 2H), 3.32 (s, 3H), 3.66 (m, 2H), 3.84 (s, 3H), 4.10 (br s, 1H), 4.57 (s, 2H), 7.30 (s, 1H). $^{13}C$ NMR (100 MHz, $CDCl_3$) *δ:* 164.8, 160.9, 138.8, 126.2, 112.6, 69.8, 58.2, 55.8, 51.3, 28.5. HR-MS: *m/z* calculated for $C_{10}H_{14}O_5$ $[M+Na]^+$: 237.0733; found: 237.0728.

**Methyl 2-isobutyl-4-methylfuran-3-carboxylate**

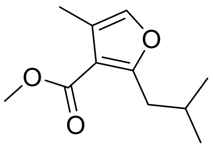

Yield: 59%. $^{1}H$ NMR (400 MHz, $CDCl_3$) *δ*: 0.91 (d, *J* = 6.5 Hz, 6H), 2.05 (non, *J* = 6.5 Hz, 1H), 2.13 (s, 3H), 2.82 (d, *J* = 7.0 Hz, 2H), 3.79 (s, 3H), 7.04 (s, 1H), $^{13}C$ NMR (100 MHz, $CDCl_3$) *δ*: 164.8, 163.2, 137.7, 120.7, 113.5, 50.9, 36.6, 28.1, 22.1, 9.8. HR-MS: *m/z* calculated for $C_{11}H_{16}O_3$ $[M+Na]^+$: 219.0997; found: 219.0997.

**Methyl 2-isobutylfuran-3-carboxylate**

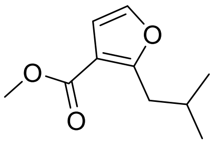

Yield: 75%. $^{1}H$ NMR (400 MHz, $CDCl_3$) *δ*: 0.93 (d, *J* = 6.5 Hz, 6H), 2.08 (non, *J* = 6.5 Hz, 1H), 2.88 (d, *J* = 7.0 Hz, 2H), 3.81 (s, 3H), 6.64 (s, 1H), 7.25 (s, H). $^{13}C$ NMR (100 MHz, $CDCl_3$) *δ*: 164.5, 162.7, 140.4, 113.5, 110.5, 51.2, 36.2, 28.3, 22.3. HR-MS: *m/z* calculated for $C_{10}H_{14}O_3$ $[M+Na]^+$: 205.0841; found: 205.0844.

***General procedure for hydrolysis***

The furan ester (1 eq.) was dissolved in a sufficient amount of 1:1 THF:$H_2O$ solution. Lithium hydroxide (2.5 eq.) was added and the reaction mixture was stirred for a minimum of overnight at either room temperature or 65 ℃. The THF was removed *in vacuo* and the remaining aqueous phase was acidified with concentrated HCl until a white precipitate formed. The precipitate was extracted with diethyl ether and the organic layer was dried over $Na_2SO_4$, filtered and concentrated *in vacuo* to afford a white or yellowish solid as the desired product.

**4-(Hydroxymethyl)-2-isobutylfuran-3-carboxylic acid (MMF1)**

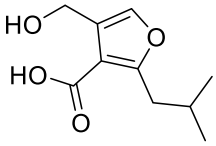

Yield: 85%. $^1$H NMR (400 MHz, $CDCl_3$) *δ*: 0.94 (d, *J* = 6.5 Hz, 6H), 2.09 (m, 1H), 2.88 (d, *J* = 7.0 Hz, 2H), 4.62 (s, 2H), 7.29 (s, 1H), 8.78 (br s, 2H). $^{13}$C NMR (100 MHz, $CDCl_3$) *δ*: 170.0, 166.1, 138.8, 125.3, 112.2, 55.6, 36.8, 28.3, 22.4. HR-MS: *m/z* calculated for $C_{10}H_{14}O_4$ $[M+Na]^+$: 221.0784; found: 221.0781.

**4-(Hydroxymethyl)-2-propylfuran-3-carboxylic acid (MMF2)**

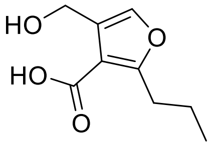

Yield: 88%. $^1$H NMR (400 MHz, $CDCl_3$) *δ*: 0.94 (t, *J* = 7.5 Hz, 3H), 1.71 (sext, *J* = 7.5 Hz, 2H), 2.97 (t, *J* = 7.5 Hz, 2H), 4.63 (s, 2H), 7.28 (s, 1H), 8.12 (br s, 2H). $^{13}$C NMR (100 MHz, $CDCl_3$) *δ*: 169.8, 166.6, 138.7, 125.3, 111.7, 55.6, 30.0, 21.3, 13.7. HR-MS: *m/z* calculated for $C_9H_{12}O_4$ $[M+Na]^+$: 207.0628, found: 207.0624.

**4-(Hydroxymethyl)-2-isopentylfuran-3-carboxylic acid (MMF3)**

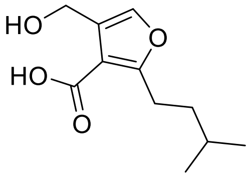

Yield: 97%. $^1$H NMR (400 MHz, $CDCl_3$) *δ*: 0.93 (d, *J* = 6.5 Hz, 6H), 1.57 (m, 3H), 3.00 (t, *J* = 7.5 Hz, 2H), 4.62 (s, 2H), 7.27 (s, 1H), 8.07 (br s, 2H). $^{13}$C NMR (100 MHz, $CDCl_3$) *δ*: 169.7, 166.9, 138.7, 125.4, 111.5, 55.6, 36.7, 27.7, 26.0, 22.3. HR-MS: *m/z* calculated for $C_{11}H_{16}O_4$ $[M+Na]^+$: 235.0941; found: 235.0942.

**2-Butyl-4-(hydroxymethyl)furan-3-carboxylic acid (MMF4)**

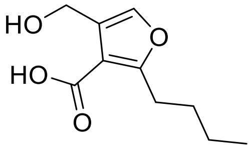

Yield: 77%. $^1$H NMR (400 MHz, $CDCl_3$) *δ*: 0.93 (t, *J* = 7.5 Hz, 3H), 1.36 (sext, *J* = 7.5 Hz, 2H), 1.68 (quin, *J* = 7.5 Hz, 2H), 3.00 (t, *J* = 7.5 Hz, 2H), 4.62 (s, 2H), 7.28 (s, 1H), 8.07 (br s, 2H). $^{13}$C NMR (100 MHz, $CDCl_3$) *δ*: 169.8, 166.8, 138.7, 125.3, 111.6, 55.6, 30.0, 27.9, 22.3, 13.7. HR-MS: *m/z* calculated for $C_{10}H_{14}O_4$ $[M+Na]^+$: 221.0784; found: 221.0781.

**4-(Hydroxymethyl)-2-pentylfuran-3-carboxylic acid (MMF5)**

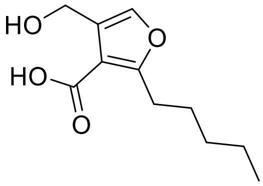

Yield: 93%. $^1$H NMR (400 MHz, $CDCl_3$) *δ*: 0.89 (t, *J* = 7.0 Hz, 3H), 1.33 (m, 4H), 1.68 (quin, *J* = 7.5 Hz, 2H), 2.99 (t, *J* = 7.5 Hz, 2H), 4.63 (s, 2H), 7.28 (s, 1H), 8.16 (br s, 2H). $^{13}$C NMR (100 MHz, $CDCl_3$) *δ*: 169.7, 166.7, 138.7, 125.3, 111.6, 55.6, 31.4, 28.1, 27.5, 22.3, 13.9. HR-MS: *m/z* calculated for $C_{11}H_{16}O_4$ $[M+Na]^+$: 235.0941; found: 235.0941.

**4-(Hydroxymethyl)-2-methylfuran-3-carboxylic acid (MMF analogue 2)**

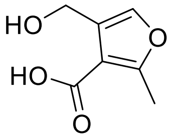

Yield: 72%. $^1$H NMR (400 MHz, $CDCl_3$) *δ*: 2.53 (s, 3H), 5.35 (s, 2H), 7.30 (s, 1H), 10.98 (br s, 2H). $^{13}$C NMR (100 MHz, $CDCl_3$) *δ*: 169.6, 163.3, 140.8, 119.2, 111.3, 60.7, 14.2. HR-MS: *m/z* calculated for $C_7H_8O_4$ $[M+Na]^+$: 179.0315; found: 179.0316.

**2-Ethyl-4-(hydroxymethyl)furan-3-carboxylic acid (MMF analogue 3)**

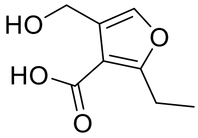

Yield: 93%. $^1$H NMR (400 MHz, $CDCl_3$) *δ*: 1.26 (t, *J* = 7.5 Hz, 3H), 3.03 (q, *J* = 7.5 Hz, 2H), 4.62 (s, 2H), 7.28 (s, 1H), 7.70 (br s, 2H). $^{13}$C NMR (100 MHz, $CDCl_3$) *δ*: 170.0, 138.7, 125.4, 111.1, 55.6, 21.8, 12.1. HR-MS: *m/z* calculated for $C_8H_{10}O_4$ $[M+Na]^+$: 193.0471; found: 193.0474.

**4-(Hydroxymethyl)-2-isopropylfuran-3-carboxylic acid (MMF analogue 4)**

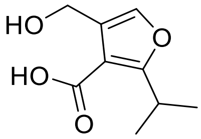

Yield: 95%. $^1$H NMR (400 MHz, $CDCl_3$) *δ*: 1.18 (d, *J* = 7.0 Hz, 6H), 3.70 (sept, *J* = 7.0 Hz, 1H), 4.54 (s, 2H), 7.20 (s, 1H), 7.95 (br s, 2H). $^{13}$C NMR (100 MHz, $CDCl_3$) *δ*: 170.6, 169.7, 138.6, 125.2, 110.2, 55.6, 27.6, 20.6. HR-MS: *m/z* calculated for $C_9H_{12}O_4$ $[M+Na]^+$: 207.0628; found: 207.0627.

**2-Hexyl-4-(hydroxymethyl)furan-3-carboxylic acid (MMF analogue 5)**

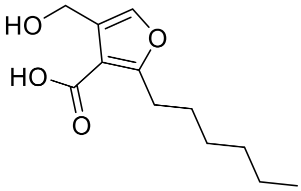

Yield: 76%. $^1$H NMR (400 MHz, $CDCl_3$) *δ*: 0.79 (m, 3H), 1.22 (m, 6H), 1.59 (m, 2H), 2.90, (t, *J* = 7.5 Hz, 2H), 4.54 (s, 2H), 7.19 (s, 1H), 8.13 (br s, 2H). $^{13}$C NMR (100 MHz, $CDCl_3$) *δ*: 169.7, 166.8, 138.7, 125.3, 111.6, 55.6, 31.5, 28.9, 28.2, 27.8, 22.5, 14.0. HR-MS: *m/z* calculated for $C_{12}H_{18}O_4$ $[M+Na]^+$: 249.1097; found: 249.1100.

**2-Heptyl-4-(hydroxymethyl)furan-3-carboxylic acid (MMF analogue 6)**

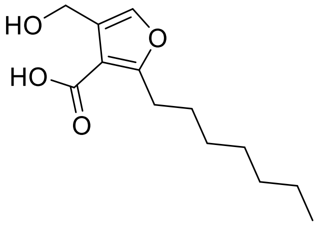

Yield: 86%. $^{1}$H NMR (400 MHz, $CDCl_3$) *δ*: 0.87 (t, *J* = 7.0 Hz, 3H), 1.28 (m, 8H), 1.67 (m, 2H), 2.98 (t, *J* = 7.5 Hz, 2H), 4.60 (s, 2H), 7.27 (s, 1H), 7.30 (br s, 2H). $^{13}$C NMR (100 MHz, $CDCl_3$) *δ*: 170.1, 167.1, 138.8, 125.5, 111.6, 55.7, 31.8, 29.3, 29.1, 28.3, 28.0, 22.7, 14.2. HR-MS: *m/z* calculated for $C_{12}H_{18}O_4$ $[M+Na]^+$: 263.1254; found: 263.1254.

**2-(But-3-enyl)-4-(hydroxymethyl)furan-3-carboxylic acid (MMF analogue 7)**

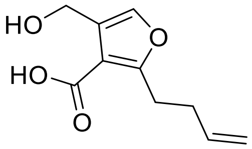

Yield: 97%. $^{1}$H NMR (400 MHz, $CDCl_3$) *δ*: 2.44 (q, *J* = 7.5 Hz, 2H), 3.10 (t, *J* = 7.5 Hz, 2H), 4.64 (s, 2H), 4.98 (dd, *J* = 10.0 and 1.5 Hz, 1H), 5.04 (dd, *J* = 17.0 and 1.5 Hz, 1H), 5.83 (dddd, *J* = 17.0, 10.0, 6.5 and 6.5 Hz, 1H), 7.29 (s, 1H), 8.27 (br s, 2H). $^{13}$C NMR (100 MHz, $CDCl_3$) *δ*: 169.5, 165.5, 138.9, 136.8, 125.3, 115.6, 112.0, 55.6, 31.8, 27.7. HR-MS: *m/z* calculated for $C_{10}H_{12}O_4$ $[M+Na]^+$: 219.0628; found: 219.0634.

**2-*sec*-Butyl-4-(hydroxymethyl)furan-3-carboxylic acid (MMF analogue 8)**

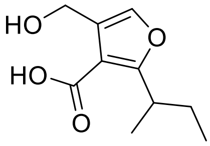

Yield: 87%. $^{1}$H NMR (400 MHz, $CDCl_3$) *δ*: 0.84 (t, *J* = 7.5 Hz, 3H), 1.25 (d, *J* = 7.5 Hz, 3H), 1.67 (quin, *J* = 7.0 Hz, 2H), 3.62 (sext, *J* = 7.0 Hz, 1H), 4.66 (s, 2H), 7.31 (s, 1H), 8.48 (br s, 2H). $^{13}$C NMR (100 MHz, $CDCl_3$) *δ*: 169.4, 169.4, 138.7, 125.2, 111.4, 55.6, 34.2, 28.3, 18.4, 11.8. HR-MS: *m/z* calculated for $C_{10}H_{14}O_4$ $[M+Na]^+$: 221.0784; found: 221.0779.

**4-(Hydroxymethyl)-2-(2-methoxyethyl)furan-3-carboxylic acid (MMF analogue 9)**

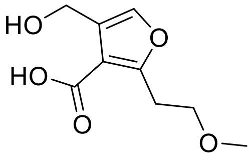

Yield: 95%. $^{1}$H NMR (400 MHz, $CDCl_3$) *δ*: 3.18 (m, 2H), 3.24 (s, 3H), 3.61 (m, 2H), 4.53 (s, 2H), 7.22 (s, 1H), 7.77 (br s, 2H). $^{13}$C NMR (100 MHz, $CDCl_3$) *δ*: 197.6, 161.9, 139.2, 125.5, 113.0, 69.9, 58.3, 55.5, 28.8. HR-MS: *m/z* calculated for $C_9H_{12}O_5$ $[M+Na]^+$: 223.0577; found: 223.0577.

**2-*iso*-Butyl-4-methylfuran-3-carboxylic acid (MMF analogue 11)**

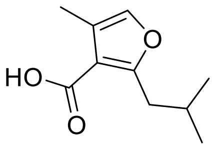

Refluxed for 3 days. Yield: 78%. $^{1}$H NMR (400 MHz, $CDCl_3$) *δ*: 0.94 (d, *J* = 6.5 Hz, 6H), 2.10 (non, *J* = 6.5 Hz, 1H), 2.18 (s, 3H), 2.82 (d, *J* = 7.0 Hz, 2H), 7.08 (s, 1H), 11.78 (br s, 1H). $^{13}$C NMR (100 MHz, $CDCl_3$) *δ*: 170.8, 165.2, 140.7, 138.1, 121.4, 113.1, 36.9, 28.3, 22.4, 10.1. HR-MS: *m/z* calculated for $C_{10}H_{14}O_3$ $[M+Na]^+$: 205.0835; found: 205.0832.

**2-*iso*-Butylfuran-3-carboxylic acid (MMF analogue 12)**

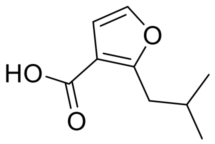

Refluxed for 3 days. Yield: 85%. $^{1}$H NMR (400 MHz, $CDCl_3$) *δ*: 0.83 (d, *J* = 6.5 Hz, 6H), 2.00 (non, *J* = 6.5 Hz, 1H), 2.80 (d, *J* = 7.0 Hz, 2H), 6.59 (s, 1H), 7.15 (s, 1H), 12.50 (br s, 1H). $^{13}$C NMR (100 MHz, $CDCl_3$) *δ*: 170.2, 164.1, 140.7, 113.3, 110.8, 36.3, 28.3, 22.3. HR-MS: *m/z* calculated for $C_9H_{12}O_3$ $[M+Na]^+$: 191.0679; found: 191.0681.

***Synthesis of MMF analogue 1***

**4-(Ethoxycarbonyl)furan-3-carboxylic acid**

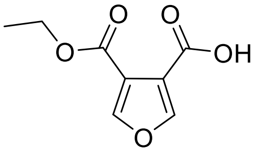

To a solution of diethyl furan-3,4-dicarboxylate (0.1 g, 0.47 mmol) in 2ml ethanol stirred at room temperature was added NaOH (0.018 g, 0.47 mmol) as a fine powder. The reaction was monitored by TLC indicating completion after 3 days. The solvent was removed *in vacuo* and the white solid dissolved in DCM was washed with 5 ml 1M $NaHCO_3$ solution. The aqueous layer was acidified with 1M HCl solution and extracted with DCM. The combined organic layers was washed with brine, dried over $MgSO_4$ and concentrated *in vacuo*. The residue was purified by flash chromatography (hexane/diethyl ether = 2:1 *v/v*) to afford 4-(ethoxycarbonyl)furan-3-carboxylic acid (0.044 g, 0.24 mmol, 51%) as white solid. $^{1}$H NMR (400 MHz, MeOD) *δ*: 1.27 (t, *J* = 7.0 Hz, 3H), 4.28 (q, *J* = 7.0 Hz, 2H), 8.14 (d, *J* = 2.0 Hz, 1H), 8.16 (d, *J* = 2.0 Hz, 1H). $^{13}$C NMR (100 MHz, MeOD) *δ*: 166.0, 164.4, 152.3, 151.6, 119.9, 118.3, 63.2, 14.4. HR-MS: *m/z* calculated for $C_8H_8O_5$ $[M+H]^+$: 185.0444; found: 185.0445.

**4-(Hydroxymethyl)furan-3-carboxylic acid (MMF analogue 1)**

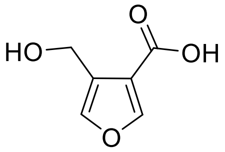

A solution of 4-(ethoxycarbonyl)furan-3-carboxylic acid (0.150 g, 0.82 mmol) in 0.7 ml dry THF was added by syringe to a stirring solution of lithium borohydride (1 M, 0.47 mmol) in THF at 0 °C under argon. The reaction mixture was allowed to warm up to room temperature while being stirred for 3 h. The reaction mixture was then cooled to 0 °C and ice cold THF-water (3 ml, 1:1) was added slowly with vigorous stirring followed by acidification with concentrated HCl. The resulting clear solution was extracted with DCM. The combined organic layers was washed with brine, dried over $MgSO_4$ and concentrated *in vacuo* to afford MMF analogue **1** (0.075 g, 0.53 mmol, 64%) as white powder. $^1H$ NMR (400MHz, MeOD) *δ*: 4.57 (s, 2H), 7.41 (s, 1H), 7.98 (s, 1H). $^{13}C$ NMR (100MHz, MeOD) *δ*: 166.9, 150.9, 142.9, 127.2, 119.2, 56.5. HR-MS: *m/z* calculated for $C_6H_6O_4$ $[M+Na]^+$: 165.0164; found: 165.0161.

***Synthesis of SCB1***

SCB1 was prepared according to modification of a reported procedure for A-factor [11]. The synthetic route for SCB1 is described in **Supplementary Figure 4**. The step involving acyl Meldrum's acid was simplified by using EDC/DMAP in one step instead of acid chloride formation and subsequent acylation in two steps. Treatment with sodium cyanoborohydride in acetic acid provided the desired double reduction product of the alkene and ketone in one step, which was followed by deprotection and column chromatography to give racemic SCB1.

**Dimethyl 2-(4-methylpentyl)malonate**

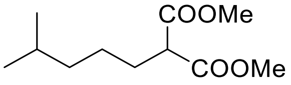

Dimethyl malonate (2.6 ml, 22.1 mmol) was slowly added to a stirring suspension of NaH (0.76 g, 30.0 mmol) in 60 ml THF at 0 ℃ and the mixture was stirred at the same temperature for 30 min. 1-Bromo-4-methylpentane (3.0 ml, 20.0 mmol) and TBAI (1.51 g, 4.0 mmol) were added and stirred at 0 ℃ for 30 min before the solution was allowed to warm to room temperature and heated to reflux overnight. The reaction mixture was concentrated *in vacuo*, acidified with 3 M HCl solution and extracted with DCM. The DCM solution was washed with water and brine, dried over anhydrous $MgSO_4$ and concentrated *in vacuo.* The residue was purified by column chromatography (petroleum ether/ethyl acetate = 10:1 *v/v*) to give dimethyl 2-(4-methylpentyl)malonate (3.54 g, 16.4 mmol, 82%) as colourless oil. $^1H$ NMR (300 MHz, $CDCl_3$) *δ*: 0.80 (d, $J$ = 6.5 Hz, 6H), 1.09-1.17 (m, 2H), 1.20-1.30 (m, 2H), 1.48 (non, $J$ = 6.5 Hz, 1H), 1.82 (q, $J$ = 8.0 Hz, 2H), 3.31 (t, $J$ = 7.5 Hz, 1H), 3.68 (s, 6H). $^{13}C$ NMR (75 MHz, $CDCl_3$) *δ*: 169.8, 52.3, 51.6, 38.3, 28.9, 27.5, 25.0, 22.3. HR-MS: *m/z* calculated for $C_{11}H_{20}O_4$ $[M+H]^+$: 217.1434; found: 217.1441.

**2-(4-Methylpentyl)malonic acid**

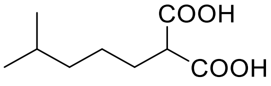

2-(4-Methylpentyl)malonate (3.54 g, 16.4 mmol) was dissolved in a mixture of KOH (9.6 g, 171.4 mmol) in 30 ml water and 60 ml methanol and the mixture was stirred overnight at room temperature. The methanol was removed *in vacuo* and the remaining aqueous solution was washed with DCM, acidified with 3M HCl solution and extracted with DCM. The DCM layers were combined, dried over anhydrous $MgSO_4$ and evaporated to yield 2-(4-methylpentyl)malonic acid (2.93 g, 15.6 mmol, 95%) as white crystals. $^1H$ NMR (300 MHz, $CDCl_3$) *δ*: 0.87 (d, $J$ = 6.5 Hz, 6H), 1.17-1.27 (m, 2H), 1.33-1.44 (m, 2H), 1.55 (non, $J$ = 6.5 Hz, 1H), 1.92 (q, $J$ = 8.0 Hz, 2H), 3.44 (t, $J$ = 7.5 Hz, 1H), 10.73 (br s, 2H). LR-MS: $[M-H]^-$ found: 187.1.

**6-Methylheptanoic acid**

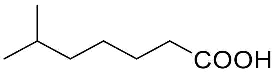

2-(4-Methylpentyl)malonic acid (2.93 g, 15.6 mmol) was dissolved in 30 ml DMF and the mixture was refluxed overnight. 50 ml 5% $NaHCO_3$ solution was added and the mixture was washed with hexane to remove the neutral impurities. The aqueous solution was acidified with 3 M HCl solution and extracted with DCM. The DCM solution was washed with brine, dried over anhydrous $MgSO_4$ and concentrated *in vacuo* to give crude product (2.2 g, quantitative) as yellow oil. $^1$H NMR (300 MHz, $CDCl_3$) *δ*: 0.85 (d, *J* = 7.0 Hz, 6H), 1.14 - 1.21 (m, 2H), 1.27 - 1.38 (m, 2H), 1.52 (non, *J* = 6.5 Hz, 1H), 1.60 (quin, *J* = 7.5 Hz, 2H), 2.34 (t, *J* = 7.5 Hz, 2H), 11.73 (br s, 1H). $^{13}$C NMR (75 MHz, $CDCl_3$) *δ*: 180.7, 38.5, 34.1, 27.8, 26.8, 24.9, 22.5. HR-MS: *m/z* calculated for $C_8H_{16}O_2$ $[M+H]^+$: 145.1223; found: 145.1222.

**2,2-Dimethyl-5-(6-methylheptanoyl)-1,3-dioxane-4,6-dione**

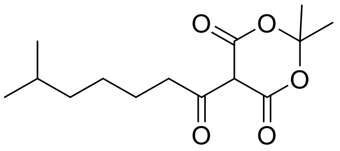

EDC·HCl (2.93 g, 15.3 mmol) was added to a solution of 6-methylheptanoic acid (2.20 g, 15.3 mmol) in 20 ml DCM and the mixture was cooled to 0 ℃. After stirring for 5 min, 4-dimethylaminopyridine (3.73 g, 30.6 mmol) dissolved in 10 ml DCM was added, followed by Meldrum's acid (2.20 g, 15.3 mmol) dissolved in 10 ml DCM. The reaction mixture was stirred under argon at room temperature for 18 h. The mixture was diluted with 20 ml DCM and washed twice with 1 M HCl, once with brine and once with water. After drying the organic phase over $MgSO_4$, the solvent was removed *in vacuo* and the residue was purified by column chromatography to yield acyl Meldrum's acid (2.0 g, 7.4 mmol, 48.4%) as yellow oil. $^1$H NMR (300 MHz, $CDCl_3$) *δ*: 0.79 (d, *J* = 6.5 Hz, 6H), 1.12-1.17 (m, 2H), 1.28-1.38 (m, 2H), 1.47 (non, *J* = 6.5 Hz, 1H), 1.61 (quin, *J* = 7.5 Hz, 2H), 1.66 (s, 6H), 2.99 (t, *J* = 7.5 Hz, 2H), 15.23 (s, 1H). $^{13}$C NMR (75 MHz, $CDCl_3$) *δ*: 170.3, 159.8, 104.4, 91.0, 38.2, 35.4, 27.5, 26.9, 26.4, 26.1, 22.2. LR-MS: $[M-H]^-$ found: 269.2.

**(1-((*tert*-Butyldimethylsilyl)oxy)-3-hydroxy propan-2-one**

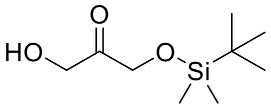

*tert*-Butyldimethylsilyl chloride (2.67 g, 17.8 mmol) dissolved in 10 ml DMF was added to a mixture of dihydroxyacetone (5 g, 55.6 mmol) and imidazole (1.51 g, 22.2 mmol) in 50 ml DMF at 0 °C. The reaction was stirred at room temperature for 17 h. 50 ml water was added and the mixture was extracted with diethyl ether. The organic phase was washed with brine, dried over anhydrous $MgSO_4$ and concentrated *in vacuo*. The residue was purified by column chromatography (petroleum ether/ethyl acetate = 5:1 v/v) to give product (1.82 g, 8.8 mmol, 50%) as colourless oil. $^1$H NMR (300 MHz, $CDCl_3$) *δ*: 0.12 (s, 6H), 0.90 (s, 9H), 2.99 (br s, 1H), 4.31 (s, 2H), 4.50 (s, 2H), further signals at 3.43-4.24 due to formation of the intermolecular ketal compounds. $^{13}$C NMR (75 MHz, $CDCl_3$) *δ*: 210.9, 25.7, 18.2, 5.6, further signals at 63.1-111.4. HR-MS: *m/z* calculated for $C_9H_{20}O_3Si$ $[M+Na]^+$ : 227.1074; found: 227.1076.

**4-(((*tert*-Butyldimethylsilyl)oxy)methyl)-3-(6-methylheptanoyl) furan-2(5H)-one**

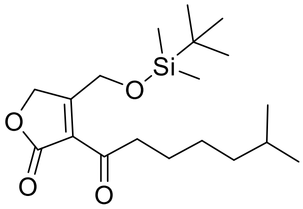

1-((tert-Butyldimethylsilyl)oxy)-3-hydroxy propan-2-one (1 eq.) and acyl Meldrum's acid (1.2 eq.) were added to 2 ml toluene already heated to 110 °C. The reaction mixture was refluxed for 3 h at which point was added 0.5 more equivalents of acyl Meldrum's acid. After stirring for another 5 h, the mixture was allowed to stand at -20 °C for 19 h and purified by column chromatography (hexanes/acetone = 60:1 *v/v*) without removal of toluene to obtain butenolide as bright yellow oil in 33% yield. $^{1}H$ NMR (300 MHz, $CDCl_3$) *δ*: 0.09 (s, 6H), 0.84 (d, *J* = 6.5 Hz, 6H), 0.89 (s, 9H), 1.10-1.36 (m, 4H), 1.47-1.63 (m, 3H), 2.96 (t, *J* = 7.5 Hz, 2H), 4.98 (s, 2H), 5.06 (s, 2H). $^{13}C$ NMR (75 MHz, $CDCl_3$) *δ*: 197.1, 181.2, 170.6, 122.4, 70.2, 61.8, 41.6, 38.7, 29.7, 27.8, 26.9, 25.7, 23.4, 22.6, -5.7. HR-MS: *m/z* calculated for $C_{19}H_{34}O_4Si$ $[M+H]^+$: 355.2299; found: 355.2306.

**4-(((*tert*-Butyldimethylsilyl)oxy)methyl)-3-(1-hydroxy-6-methyl heptyl) dihydrofuran-2(3H)-one**

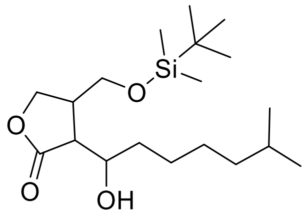

Butenolide (1 eq.) in AcOH was added dropwise to a solution of $NaBH_3CN$ (2 eq.) in AcOH at 10°C and the mixture was stirred at room temperature for 8 h. After removal of AcOH *in vacuo*, the oily residue was diluted with ethyl acetate and washed three times with 5% $NaHCO_3$ solution and once with brine. After drying the organic phase over $MgSO_4$, the solvent was removed *in vacuo* and the residue was purified by column chromatography (hexanes/ethyl acetate = 7:1 *v/v*) to yield reduction product as colourless oil in 49% yield. The reduction would result in the formation of three stereogenic centres in the molecule so NMR spectra contain a mixture of different stereoisomers. This resulted in excessive peaks and inaccurate integration. The diastereoisomers were isolated in the next step after deprotection of the silyl group. HR-MS: *m/z* calculated for $C_{19}H_{38}O_4Si$ $[M+Na]^+$: 381.2432; found: 381.2439.

**(3*R*,4*R*)-3-((*R*)-1-hydroxy-6-methylheptyl)-4-(hydroxymethyl)dihydrofuran-2(3H)-one (SCB1)**

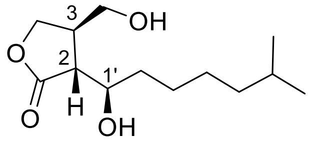

(2*R*,3*R*,1'*R*)-SCB1

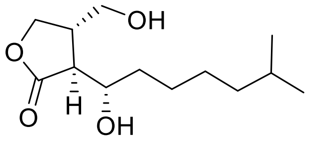

SCB1 enantiomer

TBDMS-protected butyrolactone was dissolved in a mixture of THF:HCOOH:$H_2O$ (6:3:1) and stirred overnight at room temperature. Reactions were brought up to pH 4 with saturated aqueous $NaHCO_3$ and extracted with ethyl acetate. Ethyl acetate extracts were combined, dried over $MgSO_4$ and concentrated *in vacuo*. The product was purified by flash column chromatography (hexanes/ethyl acetate = 1:1 *v/v*) to give racemic SCB1 as colourless oil in 43% yield, as well as a pair of their diastereoisomers. The NMR spectra of SCB1 enantiomers are identical to the reported data for SCB1 [12]. $^{1}H$ NMR (400 MHz, $CDCl_3$) *δ*: 0.86 (d, *J* = 6.5 Hz, 6H), 1.14-1.20 (m, 2H), 1.26-1.39 (m, 4H), 1.47-1.55 (m, 2H), 1.57-1.64 (m, 1H), 2.65 (dd, *J* = 9.5 and 4.5 Hz, 1H), 2.74-2.81 (m, 1H), 3.68 (dd, *J* = 10.5 and 6.5 Hz, 1H), 3.75 (dd, *J* = 10.5 and 5.0 Hz, 1H), 3.98 (t, *J* = 9.0 Hz, 1H), 4.01 (ddd, *J* = 9.0, 4.5 and 3.0 Hz, 1H), 4.42 (t, *J* = 8.5 Hz, 1H). $^{13}C$ NMR (100 MHz, $CDCl_3$) *δ*: 70.8, 68.3, 63.0, 49.1, 40.1, 38.8, 33.9, 27.9, 27.2, 26.1, 22.6. HR-MS: m/z calculated for $C_{13}H_{24}O_4$ $[M+Na]^+$: 267.1567; found: 267.1565.
